# Cue-guided performance is disrupted during pedunculotegmental-induced motor arrest

**DOI:** 10.64898/2026.07.09.737330

**Authors:** Madelaine C. A. Bonfils, Silas D. Larsen, Roar J. F. Sørensen, Karen Sobriel, Grace Houser, Oksana Dmytriyeva, Martin C. Tano, Hayley Burm, Rune W. Berg

## Abstract

Optogenetic stimulation of the rostral pedunculotegmental nucleus (PTg) induces global motor arrest, but it remains unclear whether this is merely a suppression of motor activity or a broader disruption of brain processes required to guide action. We developed a visuospatial cue task for rats, to test if sensory information presented during PTg-induced arrest can guide later responses. Here, we show that optogenetic stimulation during cue presentation reduces accuracy to chance level. By moving stimulation to only before or only after the cue, we found that performance was only affected when stimulation and cue presentation overlapped, that rats recover cue-guided behavior almost immediately at the end of stimulation, and that stimulation does not appear to abolish responses based on cue information acquired before arrest. These findings indicate that stimulation of the rostral PTg does not only pause motor output but transiently disrupts the ability to process and use cue information effectively.

## INTRODUCTION

Interrupting ongoing movement while remaining responsive to the environment is a fundamental feature of adaptive behavior across species. Such arrest states are typically described in motor terms, yet their behavioral consequences depend on whether sensory information received during immobility can still influence subsequent action. The pedunculotegmental nucleus (PTg) is of particular interest in this context because it has been implicated both in motor arrest and in sensorimotor processes through which sensory information can shape behavior.

The PTg is sometimes referred to as the pedunculopontine nucleus (PPN), but since it is not in the pontine region we use the former nomenclature (Watson et al. 2019). It is located in the rostral hindbrain and together with the cuneiform nucleus it forms the functionally defined mesencephalic locomotor region (MLR), an area considered essential for the generation and modulation of locomotion (Shik et al. 1966; Skinner and Garcia-Rill 1984). The anatomical boundaries of the PTg are conventionally defined by its cholinergic neurons (Rye et al. 1987; Olszewski and Baxter 1982), but the PTg contains cholinergic, glutamatergic and GABAergic populations (Luquin et al. 2018; Wang and Morales 2009; Mena-Segovia et al. 2009). Its broad ascending and descending connectivity includes motor-related structures such as the basal ganglia, motor cortex, deep cerebellar nuclei, and spinal cord (Muthusamy et al. 2007; Özkan et al. 2022), as well as regions involved in sensory processing, arousal, and cue-related behavior, including the thalamus, superior and inferior colliculi, ventral tegmental area (VTA), and sensory cortices (Özkan et al. 2022). Consistent with this organization, the PTg has been implicated in locomotion (Shik et al. 1966; Skinner and Garcia-Rill 1984; Caggiano et al. 2018; Huang et al. 2024), arousal and wakefulness (Kobayashi et al. 2004; Kroeger et al. 2017, 2022), path integration (Carvalho et al. 2020), goal-directed behavior (Thompson and Felsen 2013; Wolf et al. 2015) cue-related processing (Inagaki et al. 2022; Hormigo, Shanmugasundaram, Zhou, et al. 2021; Hormigo et al. 2019), attention (Cyr et al. 2015), reinforcement (Xiao et al. 2016; Yoo et al. 2017), and sensorimotor gating (Diederich and Koch 2005; Azzopardi et al. 2018).

Recent work has revealed functional diversity along the rostro-caudal axis of the PTg. Whereas caudal PTg stimulation is associated with initiation of locomotion and speed modulation, stimulation of the rostral PTg can produce global motor arrest (Carvalho et al. 2020; Goñi-Erro et al. 2023; Kaur et al. 2025). While stimulation of Vsx2-positive glutamatergic neurons in the rostral PTg (also known as Chox10+) is sufficient to induce global motor arrest (Goñi-Erro et al. 2023), perturbations of PTg neurons or projections have been shown to affect movement direction (Tello et al. 2024), locomotor speed (Roseberry et al. 2016; Xiao et al. 2016), and ongoing purposive behaviors (Gut et al. 2022). Together, these findings suggest that the behavioral consequences of PTg activation are not limited to locomotor output, and may depend on anatomical location, cell type, projection target, and ongoing behavioral context.

However, it remains unclear if PTg-induced motor arrest reflects a selective interruption of motor output or if it also disrupts sensorimotor processes required to transform new sensory information into action. This distinction cannot easily be resolved from spontaneous behavior alone, because animals may resume ongoing actions after stimulation without necessarily having processed information presented during arrest.

To address this, we developed a self-paced visuospatial cue task, which temporally separates cue presentation from response execution, to determine if visual cue information presented during PTg-induced arrest can guide later behavior. When rostral PTg stimulation overlapped with cue presentation, accuracy decreased to chance-level and was not rescued by extending cue presentation. This impairment was temporally specific: accuracy was largely preserved when stimulation ended before cue presentation or when stimulation began after cue offset. Because stimulation was delivered unilaterally, we also asked whether impaired performance was influenced by a lateralized response bias. Side bias emerged on stimulation trials when accuracy was impaired but was not systematically related to the stimulated hemisphere. These findings indicate that rostral PTg stimulation disrupts cue-related processing, in addition to suppressing motor output.

## RESULTS

### Viral expression and motor arrest

Eleven rats received bilateral injections of an AAV2/9-CamKIIa-ChrimsonR-mScarlet virus for opsin-expression and optic fiber implants targeting the rostral PTg after reaching training criteria on the visuospatial cue task (VSCT) (Materials & Methods: Training protocol). Four weeks later, ten animals displayed clear global motor arrest during optogenetic stimulation (10 ms pulses at 50 Hz). Postmortem histology confirmed viral expression and fiber placement in the rostral PTg by either light-sheet fluorescence microscopy-based 3D volumetric imaging of cleared, ChAT-immunostained hemispheres (**Figure 1A**) or four-channel multiplex RNAscope fluorescence in situ hybridization on sagittal sections including a ChAT probe (**Figures 1B and 1E**).

**Figure 1.**
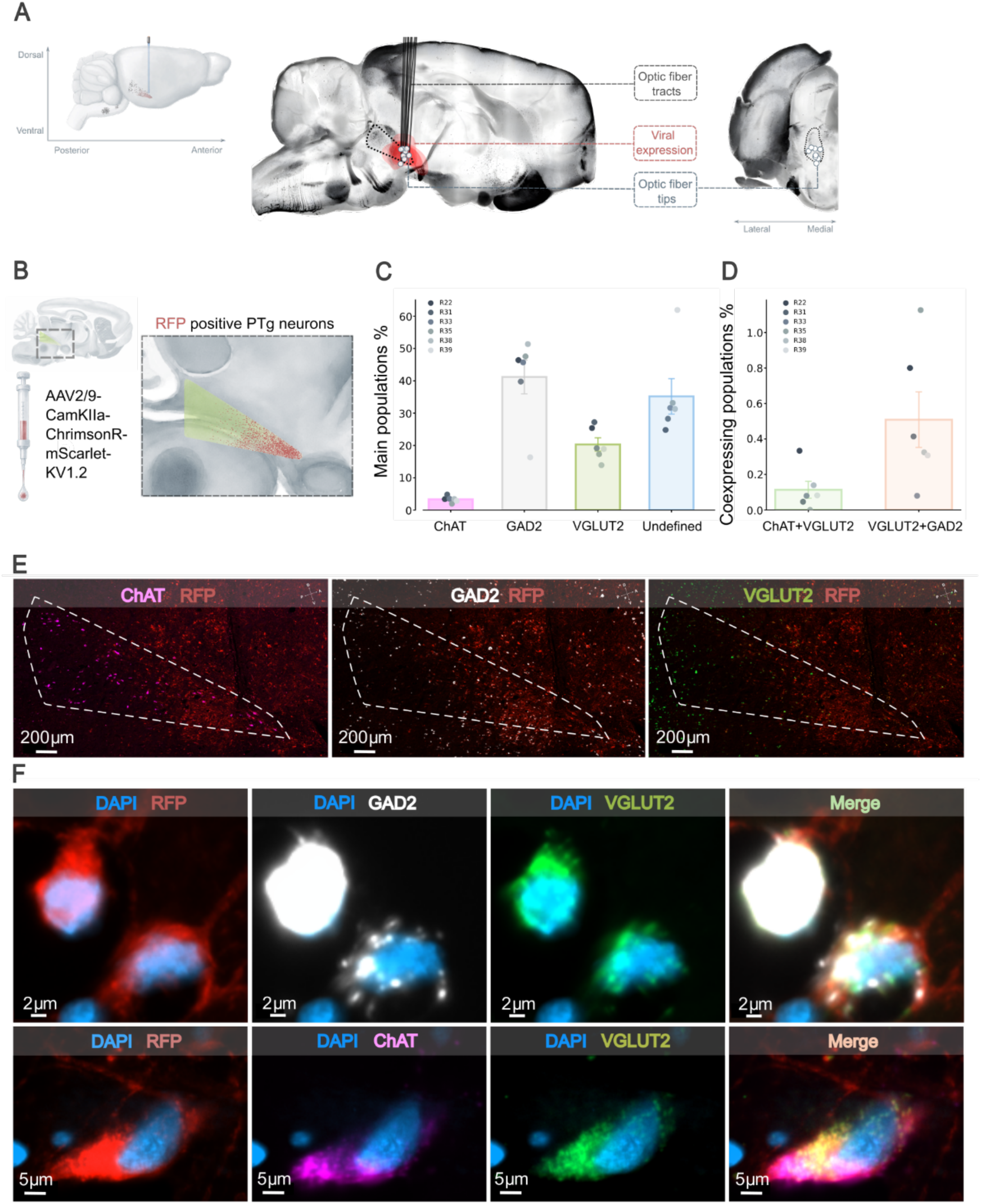
Fiber placement, viral expression and cell types. A) Composite illustration of viral distributions (red), fiber tracts (dark grey lines), and fiber tips (circles) from 3D cleared tissue. Tissues were aligned to a ChAT-stained reference rat brain. B) Schematic of a sagittal section of rat brain with the PTg (green) and CamKIIa expressing population (red) that was analyzed using multiplex RNAscope. The analysis was limited to virally transduced cells within the PTg. The boundary of the PTg was defined by its ChAT positive neurons and viral signal intensity was enhanced with RFP. C) Estimated fractions of neuronal populations found within the PTg (14189 cells in total, n = 6) based on expression levels of ChAT, GAD2 and VGLUT2. D) Estimated fractions of neuronal cell populations co-expressing GAD2/VGLUT2 or ChAT/VGLUT2. E) Photomicrographs of 12 µm sagittal sections of the PTg: Left, co-expression of ChAT (magenta) and RFP (red); Center, co-expression of GAD2 (white) and RFP (red); Right, co-expression of VGLUT2 (green) and RFP (red). F) Top: Photomicrograph of two cells co-expressing GAD2 (white) and VGLUT2 (green). Bottom: Photomicrograph of a cell co-expressing ChAT (magenta) and VGLUT2 (green). Schematic diagram of sagittal section adapted with permission from Paxinos G and Watson C (The Rat Brain in Stereotaxic Coordinates, 7th Edition, 2014). Blue = DAPI; Red = RFP. Chat = Choline acetyltransferase, GAD2 (Gad65) = Glutamic Acid Decarboxylase 2, VGLUT2 = Vesicular glutamate transporter 2, RFP = red fluorescent protein.

To characterize the transduced population, we combined multiplex RNAscope with RFP immunolabeling within the ChAT-defined PTg boundary (**Figure 1B–F**). Viral expression was distributed across multiple neuronal populations, including GAD2-positive (41.18%), VGLUT2-positive (20.31%), and ChAT-positive cells (3.32%), while 35.18% of virally expressing cells were unclassified (**Figures 1C and 1E**). Rare co-expression was observed for GAD2/VGLUT2 (0.51%) and ChAT/VGLUT2 (0.11%) cells (**Figure 1D and 1F**). Thus, stimulation was targeted to the rostral PTg but was not restricted to a single molecularly defined cell type.

### PTg stimulation during cue presentation disrupts performance

The visuospatial cue task was designed to test cue-related processing during stimulation-induced motor arrest. In each trial, the rat initiated the trial by nose-poking a small white rectangle at the bottom of the touchscreen, which encouraged the rat to orient toward the screen before cue presentation. A visual cue (white plus sign) then appeared in one of three possible screen positions: left, center, or right. After the cue disappeared, three visually identical stimuli appeared simultaneously in the same three positions. To receive a reward, the rat had to nose-poke the stimulus located in the same position as the preceding cue (**Figure 2C and 2D; Video S1**).

**Figure 2.**
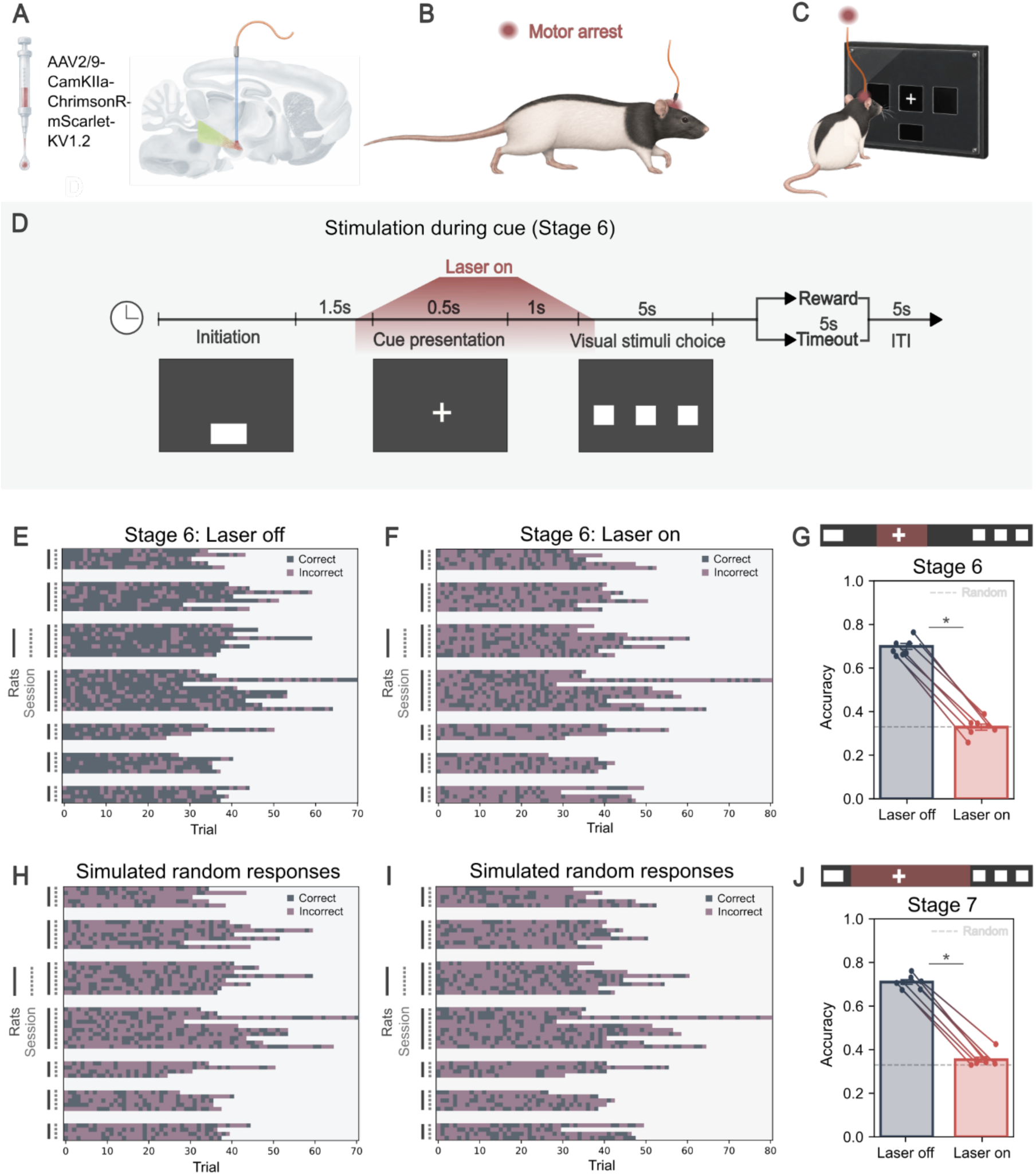
Performance is obstructed when PTg stimulation occurs during cue presentation. A) Virus injection into the rostral PTg (green triangle), and location of the implanted optical fiber in a sagittal section. Optogenetics was performed unilaterally. B) Illustration of a rat pose during motor arrest. C) Rat facing the touchscreen receiving optogenetic stimulation during cue presentation. D) Stimulation during cue (Stage 6) overview: Trial initiation with a nose poke (first screen). A cue (plus sign) is then presented in one of three positions (second screen). Three identical visual stimuli appear (third screen), the rat selects one, but only the choice of the same stimulus position as the preceding cue is rewarded, while incorrect responses result in a timeout. E-F) Performance in “Laser oe” (E) and “Laser on” trials (F). Each vertical bar corresponds to one rat, each horizontal line is a session with dots indicating a correct (dark blue) or incorrect (purple) trial outcome. G) Accuracy on “Laser oe” (blue) vs “Laser on” (red) trials. Chance level indicated (dashed line). Mean ± SEM = -0.37 ± 0.02, n = 7, p = 0.016. H-I) Simulated data with random accuracy (33%) for comparison (E, H) and (F, I). J) Accuracy during “Laser oe” (blue) and “Laser on” (red) trials on Extended stimulation sessions (Stage 7; 4.5 s or 5.5 s) and cue presentation (2.5 s or 3.5 s). Mean ± SEM = -0.35 ± 0.02, n = 7, p = 0.016. Wilcoxon signed-rank two-sided test (“Laser oe” vs “Laser on”. ITI = Intertrial Interval. Schematic diagram of sagittal section adapted with permission from Paxinos G and Watson C (The Rat Brain in Stereotaxic Coordinates, 7th Edition, 2014).

The main stimulation condition, Stage 6 (**Video S2**), tested performance when cue presentation overlapped with motor arrest. PTg was stimulated for 2.5 s on 50% of trials in a pseudorandom order. Stimulation began 0.5 s before cue presentation and continued until 0.5 s after the three response stimuli appeared. Thus, on stimulation trials, the cue was presented only while stimulation was active, and rats were required to respond after stimulation ended. Because stimulation and control trials were pseudorandomly interleaved within each session, laser-off trials served as the primary within-subject control condition for behavioral analyses. Seven of the ten rats that displayed motor arrest met the task’s performance criteria, and unless otherwise stated, each analyzed stimulation stage included at least four sessions per rat, with each session containing ≥50 completed choice trials after excluding premature responses and omissions.

The first question was whether rats could perform the task when cue presentation overlapped with stimulation-induced motor arrest. Stimulation of the rostral PTg during cue presentation significantly decreased accuracy by 37 ± 2 percentage points, from 70 ± 1% on laser-off trials to 33 ± 1% on laser-on trials (Wilcoxon signed-rank test, n = 7, p = 0.016; **Figure 2G**). Because this was a three-choice task, 33% accuracy represents chance-level performance. Accuracy on laser-off trials was significantly above chance, whereas accuracy on laser-on trials was indistinguishable from chance (Wilcoxon signed-rank tests against 33% chance level: laser off, p = 0.016; laser on, p = 1.000). To provide a trial-level visual reference for chance performance, we simulated datasets matched to the experimental data in session and trial number, but with randomly assigned outcomes corresponding to 33% expected accuracy (**Figure 2E, 2F, 2H, and 2I**).

To test if this decrease in accuracy could be explained by a transient reaction to stimulation onset, such as a brief shift in attention or the experience of being arrested, rats were tested in an extended cue-presentation condition (**Stage 7, Figure 2J, Video S3**). Two versions of Stage 7 were used: 2.5 s cue presentation with 4.5 s stimulation, and 3.5 s cue presentation with 5.5 s stimulation. Stimulation began before cue presentation and continued until after the response stimuli appeared, as in Stage 6. Since we did not observe a difference between the two Stage 7 versions, the datasets were merged for the final analysis. Accuracy decreased by 35 ± 2 percentage points, from 71 ± 1% on laser-off trials to 35 ± 1% on laser-on trials (Wilcoxon signed-rank test, n = 7, p = 0.016; Figure 2J). While we cannot rule out that the onset of stimulation affected the animals’ experience of the trial, the persistence of near-chance performance despite prolonged cue presentation suggests that the deficit was not caused solely by a brief distraction or a transient attentional shift at stimulation onset.

### Off-cue stimulation has no effect

To determine whether stimulation-induced motor arrest produced lingering effects that could impair performance after stimulation ended, stimulation was delivered before cue presentation in Stage 8, during the first black-screen interval between trial initiation and cue onset (**Figure 3A, Video S3**). Two versions of Stage 8 were used: either 2.2 s stimulation ending 0.3 s before cue presentation, or 2.5 s stimulation ending exactly at the onset of cue presentation. The first black-screen interval was extended to 4 s on stimulation trials only, while laser-off trials remained identical to those in Stages 6 and 7. Because response time and accuracy did not differ detectably between the two Stage 8 versions, the datasets were merged for the final analysis.

**Figure 3.**
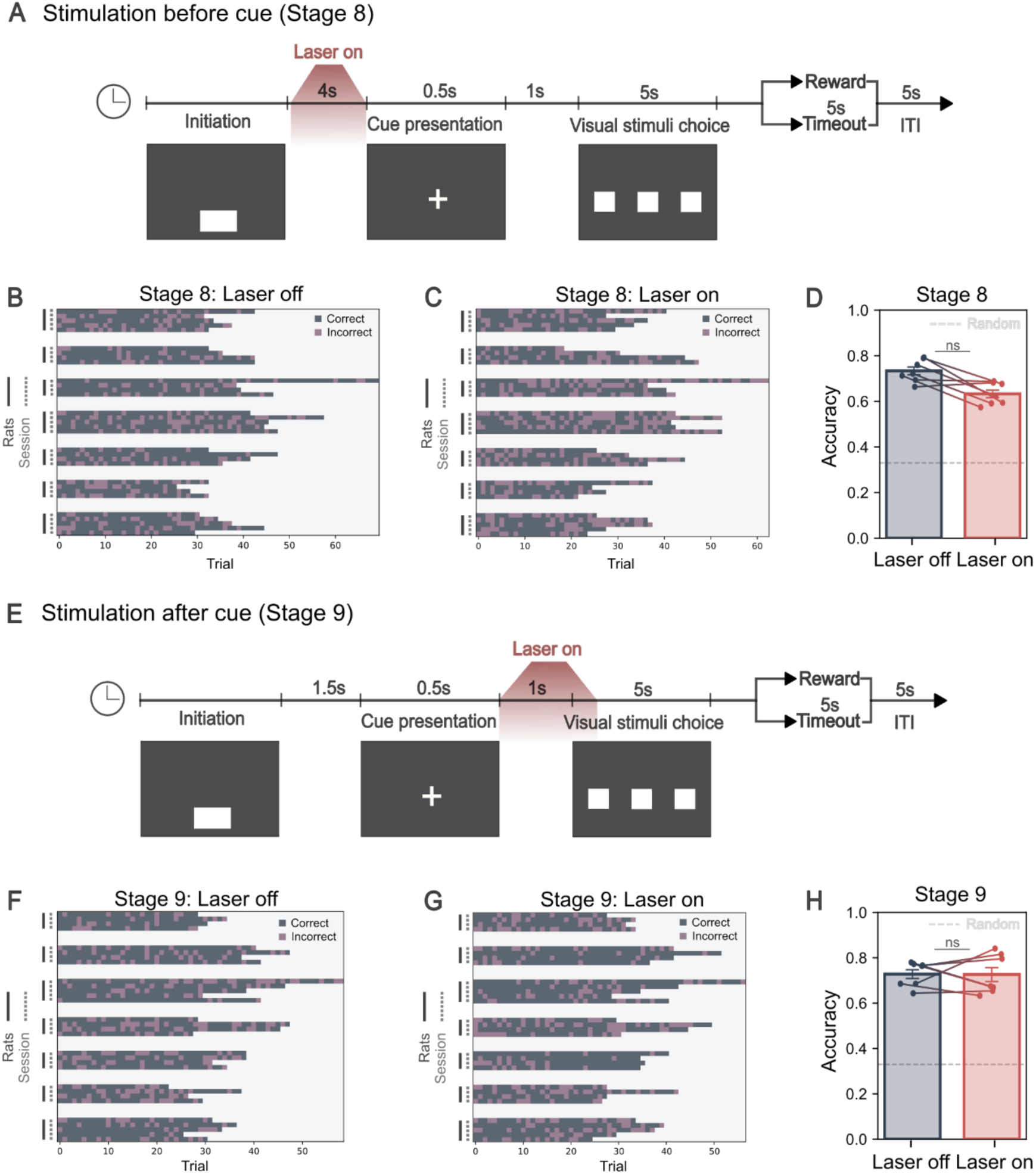
Oe-cue stimulation has no significant eeect on performance. A) Stimulation before cue (Stage 8) trial overview. Stimulation began 0.5 s after trial initiation and ended either 0.3 s before or exactly at the onset of cue presentation. The first black screen interval was increased to 4.0 s during stimulation trials. B-C) Stage 8 “Laser oe” (B) and “Laser on” (C) trials. Each vertical bar corresponds to one rat, each horizontal line is a session with dots indicating a correct (dark blue) or incorrect (purple) trial outcome. D) Dieerence between accuracy on “Laser oe” (blue) and “Laser on” (red) trials is not significant when stimulation occurs before the cue. Mean ± SEM = -0.09 ± 0.03, n = 7, p = 0.078. Chance level indicated (dashed line). E) Stimulation after cue (Stage 9) trial overview. Stimulation occurred either 0.2 or 0.5 s after cue presentation ended. F-G) Stage 9 “Laser oe” (F) and “Laser on” (G) trials. H) Dieerence between accuracy on “Laser oe” (blue) and “Laser on” (red) trials is not significant when stimulation occurs after cue presentation. Mean ± SEM - 0.00 ± 0.03, n = 7, p = 0.938. Wilcoxon signed-rank two-sided test (Laser oe vs Laser on).

While accuracy was reduced by 9 ± 3 percentage points on laser-on trials, this effect was not significant (Wilcoxon signed-rank test, n = 7, p = 0.078; **Figure 3B–D**), and response time did not differ between laser-off and laser-on trials (laser off: 1.73 ± 0.03 s, laser on: 1.73 ± 0.05 s, Wilcoxon signed-rank test, n = 7, p = 0.947). These findings suggest that the behavioral effects of rostral PTg stimulation do not persist long enough to significantly impair cue-related performance after stimulation ends.

### Performance is preserved during post-cue stimulation

In the visuospatial cue task, rats had to withhold responding during cue presentation. Interaction with the screen within this window was recorded as premature responses and led to termination of the trial, an error tone, and was followed by a 5 s timeout. Thus, the rat must receive the information of the cue location but withhold its response until after cue offset. This leaves an interval where the rat has seen the cue but has yet to complete its response. In Stage 9, we test if cue-guided responses were impaired if briefly interrupted by stimulation of the rostral PTg *after* cue presentation.

Rats were tested on two versions of Stage 9 with 2.5 s stimulation starting either 0.2 s or 0.5 s after the cue presentation ended (**Figure 3E**). Since accuracy did not differ significantly between the two versions, the datasets were merged for the final analysis. When stimulation followed cue presentation, accuracy was unchanged between Laser off and Laser on trials (difference: 0±3 percentage points, Wilcoxon signed-rank test, n = 7, p = 0.938; Figure 3F–H). This contrasts with the marked impairment observed when stimulation overlapped with cue presentation.

One caveat is that on some trials, rats oriented rapidly after cue offset and were already aligned with the cued response location by the time stimulation began. These trials may reduce Stage 9’s sensitivity to detect if stimulation disrupts the maintenance or use of cue information. Nevertheless, these findings suggest that rostral PTg stimulation does not abolish the ability to use cue information acquired before stimulation onset, though a more subtle impairment cannot be excluded.

### Side preference emerged when performance was impaired

A reduction in accuracy could reflect a non-cue biased response preference, in which the animal chooses a stimulus position more often than expected by chance, i.e. a “side bias”. Therefore, we examined if stimulation altered side bias across task stages (**Figure 4**). On “Laser-off” trials, responses were distributed approximately evenly among the three response positions in each stage, and when summarized across task stages (Left: 0.31 ± 0.021 SEM; Center: 0.36 ± 0.02 SEM; Right: 0.32 ± 0.01 SEM; Friedman chi-square test; n = 7, p = 0.156) (**Figure 4C_1_-F_1_**). Thus, we found no strong evidence for a baseline side bias in the absence of stimulation.

**Figure 4.**
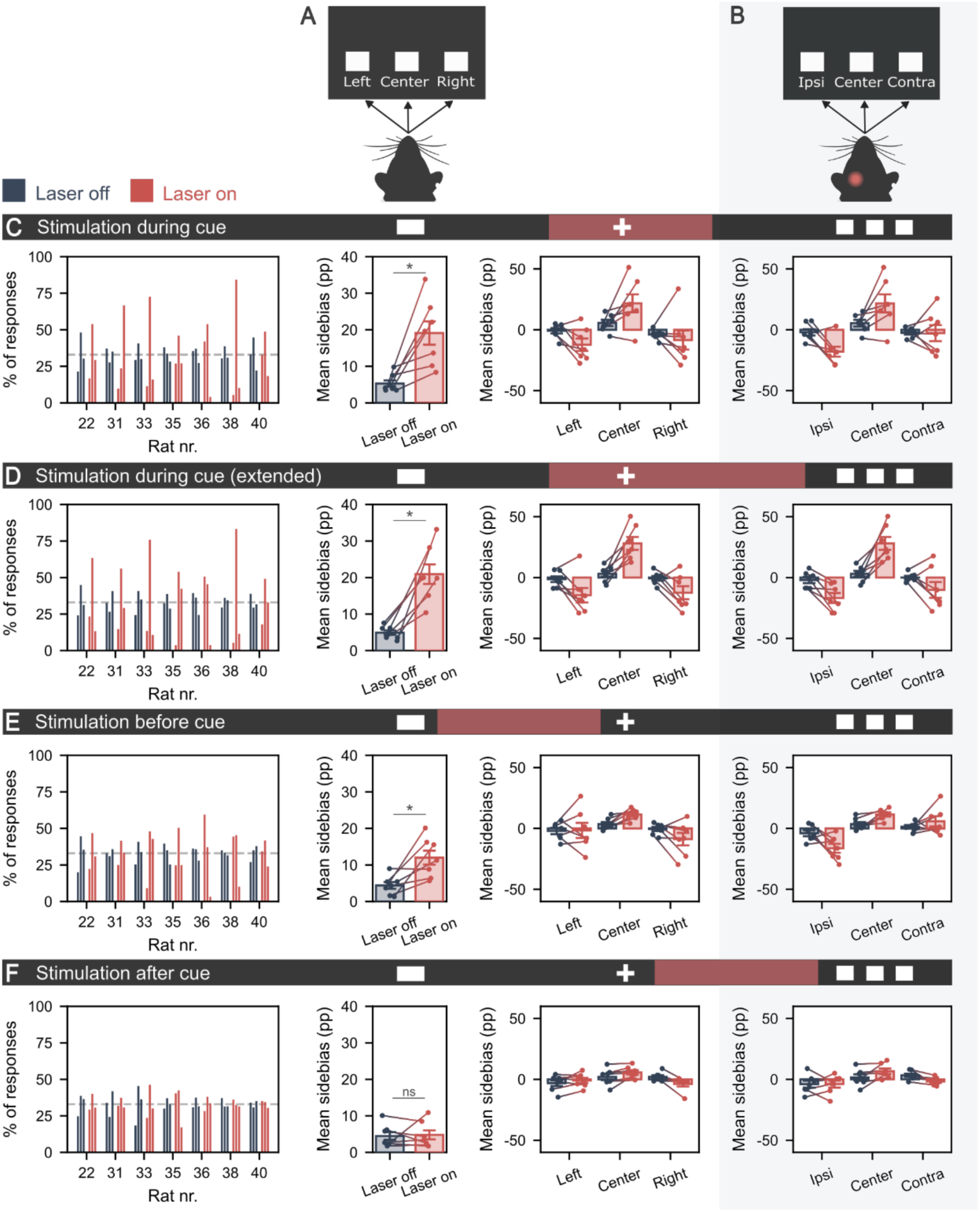
Testing for side bias across stimulation stages. A) The three choices on the touch screen: Left, center and right. B) Definitions of unilateral stimulation vs side bias: Ipsi = ipsilateral refers to responses to the same side as stimulation, while Contra = contralateral are responses made to the opposite side of stimulation. Center responses were not assigned as ipsilateral or contralateral. For each stimulation stage (C-F), left panel (panel 1): percentage of responses from individual rats to each position for “Laser oe” and “Laser on” trials; second left panel (panel 2): mean side bias (percentage points, PP) of “Laser oe” vs. “Laser on” trials; panel 3: mean PP changes for each position of “Laser oe” and “Laser on” trials; and right panel (panel 4): mean PP changes on the ipsi/contra sides of “Laser oe” and “Laser on” trials. C) Stage 6: Stimulation during cue. C1) Response distribution per rat (Laser oe/on). C2) Side bias (PP): 13.82 ± 3.96, n = 7, p = 0.016. C3) Position specific PP change (n = 7, p = 0.012): Left = -11.44±4.83; Center =16.06 ± 6.54; Right = -4.63 ± 6.94. C4) Ipsi/contra PP change: Ipsi -15.89 ± 4.45 vs Contra -0.33±6.24, n=7, p=0.109. D) Stage 7: Extended stimulation (Laser 4.5 or 5.5 s; Cue 2.5 or 3.5 s). D1) Response distribution per rat (Laser oe/on). D2) Side bias (PP): 16.00 ± 3.02, n = 7, p = 0.016. D3) Position specific PP change (n = 7, p = 0.021): Left = -13.14 ± 5.36; Center =24.92 ± 4.96; Right = -11.78 ± 5.36. D4) Ipsilateral vs. Contralateral PP change: Ipsi -14.77 ± 3.66 vs Contra -10.15 ± 6.06, n = 7, p = 0.938. E) Stage 8: Stimulation before cue. E1) Response distribution per rat (Laser oe/on). E2) Side bias (PP): 7.61 ± 2.39, n = 7, p = 0.031. E3) Position specific PP change (n = 7, p = 0.276): Left = -0.43 ± 5.41; Center = 7.90 ± 1.92; Right = -8.33 ± 4.58. E4) Ipsilateral vs. Contralateral PP change: Ipsi -12.34 ± 4.51 vs Contra -4.44 ± 4.29, n = 7, p = 0.109. F) Stage 9: Stimulation after cue. F1) Response distribution per rat (Laser oe/on). F2) Side bias (PP): 0.32 ± 1.47, n = 7, p = 0.688. F3) Position specific PP change (n = 7, p = 0.156): Left = -2.22 ± 1.80; Center = 3.70 ± 1.70; Right = -5.92 ± 2.31. F4) Ipsilateral vs. Contralateral PP change: Ipsi 0.07 ± 2.95 vs Contra -5.12 ± 1.54, n = 7, p = 0.297. Chance level indicated (dashed line). Friedman chi squared test: Panel 3, Wilcoxon signed-rank test: Panel 2 and Panel 4.

However, on laser-on trials, side bias increased significantly in Stage 6 (increase: 13.82 ± 3.96 percentage points, Wilcoxon signed-rank test, n = 7, p = 0.016), Stage 7 (increase: 16.00 ± 3.02 percentage points; n = 7, p = 0.016), and Stage 8 (increase: 7.61 ± 2.39 percentage points; n = 7, p = 0.031), but not in Stage 9 (increase: 0.32 ± 1.47 percentage points; n = 7, p = 0.688, **Figure 4C_2_- F_2_**). In Stages 6–8, this bias appeared mainly as an increase in responses to the center stimulus, with corresponding decreases in responses to the left and right stimuli, although this position-specific trend was not significant (**Figure 4C_3_-E_3_**). Across stages, larger impairments in accuracy were accompanied by larger increases in side bias. Side bias increased most strongly in Stages 6 and 7, where accuracy was reduced by 37 and 35 percentage points, respectively. A smaller increase in side bias was observed in Stage 8, where accuracy was reduced by 9 percentage points, whereas no side bias emerged in Stage 9, where accuracy was unchanged. This pattern suggests that side bias was greatest when rats were least able to use cue information to guide their responses.

To test whether stimulation biased responses toward or away from the side of stimulation, we compared lateral responses relative to the stimulated hemisphere. Ipsilateral responses were responses to the stimulus on the same side as stimulation, whereas contralateral responses were responses to the stimulus on the opposite side. Because center responses could not be assigned to either category, this analysis included only responses to the left and right stimuli. We found no significant ipsilateral or contralateral response bias in any stage (**Figure 4B_4_-F_4_**).

Together, these results indicate that rostral PTg stimulation did not produce side bias as a direct function of stimulation. Instead, side bias emerged when cue-guided performance was impaired, consistent with animals relying more on a default or biased response strategy when cue information could not be effectively used. Overall, the behavioral results show that rostral PTg stimulation strongly disrupts performance when it overlaps with cue presentation, whereas performance is largely preserved when stimulation occurs before or after cue presentation.

## DISCUSSION

It was recently discovered that stimulation of the brainstem nucleus PTg induces a complete arrest of movement. This is a striking and puzzling phenomenon to observe: the animal appears frozen, yet shows no signs of fear or distress, and resumes its prior behavior the instant stimulation ends, as though nothing had happened. Is the arrest purely a motor phenomenon? Or does it also suspend other brain processes, such as sensory processing, decision-making and planning of goal-directed behavior? Is the animal aware of its surroundings but simply unable to move, or is brain function more broadly disrupted during the arrest?

Here, we demonstrate that PTg-induced motor arrest during the presentation of a visual cue abolishes cue-guided task performance, reducing accuracy to chance. This impairment is temporally specific to the overlap between cue presentation and optogenetic stimulation. When the stimulation does not overlap with cue presentation, occurring either before or after the cue, performance is intact. This temporal specificity suggests that PTg-induced motor arrest is not merely a suppression of motor output but also interferes with processes required to use sensory cue information to guide later responses. In parallel with the decline in accuracy, a response bias emerged on stimulation trials, suggesting that animals relied more on biased responding when cue-guided performance was disrupted.

Given that subsets of PTg neurons have been shown to induce ipsilateral orienting (Hormigo, Zhou, and Castro-Alamancos 2021) and turning (Tello et al. 2024), the emergence of side bias during stimulation suggests an orienting component to the behavioral deficit. However, we found no evidence that such mechanisms account for the side bias observed here, as responses were not systematically biased toward the ipsilateral stimulus relative to the stimulation side. This analysis was limited by the inability to unambiguously assign responses to the center stimulus as ipsilateral or contralateral, and by the lack of control over head position at stimulation onset. Importantly, if stimulation induced an ipsilateral orienting bias, similar changes in side bias would be expected across all stages. Instead, side bias increased as accuracy declined and was absent when performance was unaffected, as observed when stimulation occurred after cue presentation in Stage 9.

### Ready, Set, Pause!

Motor arrest can occur as part of several distinct processes, including defensive freezing, cancellation of ongoing action, or a transient pause that allows behavior to resume. Freezing behavior, classically defined as a state of attentive immobility in response to a threat (Kozlowska et al. 2015), is unlikely to explain the arrest induced by PTg stimulation as neither we nor other laboratories investigating PTg-induced motor arrest have observed additional behavioral features characteristic of freezing behavior (Goñi-Erro et al. 2023; Kaur et al. 2025; Huang et al. 2024). Instead, optogenetic stimulation of the PTg resembles that of a pause-and-play phenotype, characterized by rapid onset and offset of arrest with immobility that preserves the animal’s posture at stimulation onset (Goñi-Erro et al. 2023). Conversely, stimulation of the ventrolateral periaqueductal grey (vlPAG), a region associated with defensive freezing, produced prolonged immobility, with stereotyped arrest postures, and failed to resume pre-stimulation behavior (Goñi-Erro et al. 2023), whereas animals stimulated in the rostral PTg would often resume behaviors such as walking (Goñi-Erro et al. 2023), eating, or swimming (Huang et al. 2024) at stimulation offset. Thus, in contrast to action cancellation, which would require re-initiating behavior after stimulation, PTg stimulation can produce a reversible interruption of movement that preserves the behavioral state and allows animals to resume ongoing actions immediately upon stimulation offset. Consistent with this interpretation, we observed no increase in response latency or omission rate when stimulation was delivered before the cue (data not shown), as would be expected if animals had to re-establish task engagement after a cancelled action. Stimulating after the cue (Stage 9) provides a less direct test of action cancellation than the before-cue condition, because stimulation began only after a short post-cue interval. This delay was intended to allow cue information to guide response selection before stimulation onset, but it also allowed some animals to approach the selected response location before stimulation began. As a result, some laser-on trials may have required only minimal additional movement after stimulation ended. Even so, if stimulation had reset task engagement, cancelled an already selected response, or abolished access to cue information acquired before the onset of stimulation, we would expect detectable behavioral consequences, such as increased omissions or reduced accuracy. The absence of such effects therefore argues against action cancellation as the primary explanation for PTg-induced motor arrest in this task.

This interpretation aligns with proposed fast-acting pause networks involving the PTg, subthalamic nucleus (STN), and the substantia nigra *pars reticulata* (SNr), which transiently delay motor output while determining whether to continue or cancel an action (Schmidt et al. 2013; Mosher et al. 2021). Importantly, describing the effect as a transient pause does not by itself explain the collapse in cue-guided performance. We therefore propose that stimulation of the rostral PTg interferes with cue-related processing in addition to pausing motor output.

### Goal-directed behavior and cue-related processing in the PTg

In our visuospatial cue task, rats continuously scan the screen while awaiting a cue, whose pseudorandom position prevents its prediction. Thus, prior to cue presentation, animals are prepared to act but have not yet selected a movement direction. A successful trial requires rapid processing of the visual cue and integration of this information with motor-control circuits to select and initiate the appropriate response. The finding that performance was abolished only when stimulation overlaps with cue presentation suggests disruption of cue-related processes or early goal-directed behavior rather than response execution or reinforcement alone.

This interpretation is consistent with a growing literature implicating the PTg in both cue responsiveness (Gut et al. 2022; Wolf et al. 2015; Inagaki et al. 2022; Hormigo et al. 2019; Pan and Hyland 2005) and action initiation (Gut et al. 2022; Thompson and Felsen 2013; Tello et al. 2024; Masini and Kiehn 2022). In line with this, the PTg has been shown to respond to both visual and auditory stimuli (Huang et al. 2024; Noftz et al. 2020; Yau et al. 2016; Lee et al. 2014; Pan and Hyland 2005), with selectivity toward salient stimuli (Hormigo, Shanmugasundaram, Zhou, et al. 2021) and learned cues (Inagaki et al. 2022).

In both signaled avoidance (Hormigo et al. 2019) and delayed response tasks (Inagaki et al. 2022), perturbation of the PTg or closely related regions (midbrain reticular nucleus = MRN), was shown to block cue-evoked responses despite preserved motor capability. Specifically, both inhibition of the PTg/MRN (Inagaki et al. 2022), as well as selective stimulation of GABAergic PTg neurons (Hormigo et al. 2019), blocks “go-cue” responses, while non-selective stimulation of the PTg/MRN can mimic cue-driven responses, directly inducing the cued behavior even in the absence of the external signal (Inagaki et al. 2022). In these paradigms, however, the relevant cue primarily signals when to release a prepared or learned action. By contrast, in our task the cue specifies which spatial response should be selected. Our findings therefore extend this role by showing that rostral PTg stimulation can also disrupt behavior when newly presented cue information is needed to determine the appropriate later action.

Our task temporally separated cue presentation from response execution but we cannot identify which specific step was disrupted: cue detection, maintenance of cue information, action selection, or initiation of the selected response. Related work provides converging evidence that PTg-dependent circuits are especially important during early stages of action initiation. In a self-paced lever-press task, Gut et al. (2022) showed that perturbation of GABAergic efferents from the PTg to the SNc, after action initiation, pauses behavior without degrading accuracy, whereas perturbation during initiation increases failure rates significantly (Gut et al. 2022). This aligns with our observation that stimulation after cue presentation preserves performance, while stimulation during cue presentation abolishes it, raising the possibility that rostral PTg stimulation disrupted an early cue-dependent action-selection or initiation process rather than sensory cue processing per se. However, the deficit we observed was not a partial reduction in performance: when stimulation overlapped with cue presentation, accuracy fell to chance. Together, these findings support a mechanism in which PTg activity contributes to early cue-dependent stages required to transform cue information into a correct action, while ongoing or already specified actions can be paused and resumed.

Although the precise connectivity underlying these behaviors requires further investigation, the PTg projects to multiple regions involved in sensory processing and action control, including the basal ganglia, thalamus, and superior colliculus, which contribute to goal-directed behavior, orienting movements, and action selection (Wolf et al. 2015; Thompson and Felsen 2013; Inagaki et al. 2022). Thus, we propose that rostral PTg stimulation pauses motor output, likely through a subset of glutamatergic neurons (Huang et al. 2024; Goñi-Erro et al. 2023), while also disrupting processes that allow newly presented cue information to guide subsequent behavior. While we cannot pinpoint the exact computational step affected, this role is consistent with the PTg’s involvement in sensorimotor gating (Diederich and Koch 2005), potentially regulating the flow of sensory information to downstream motor circuits, perhaps in part through GABAergic projections to basal ganglia and thalamus.

### Cellular diversity and behavioral consequences of PTg perturbation

The involvement of the PTg in both motor arrest and cue-related processing raises the possibility that the behavioral phenotype observed here reflects recruitment of multiple neuronal populations or projection pathways. Stimulation of glutamatergic neurons in the rostral PTg is sufficient to induce global motor arrest (Goñi-Erro et al. 2023; Huang et al. 2024). However, non-selective stimulation of the PTg may engage additional behavioral processes. For example, selective stimulation of Vsx2-positive glutamatergic neurons induces motor arrest without inducing either place preference or aversion (Goñi-Erro et al. 2023), whereas optogenetic stimulation of cholinergic PTg neurons induces place preference, and inhibition induces place aversion (Xiao et al. 2016). Moreover, stimulating glutamatergic efferents from the PTg to the VTA promoted a preference for stimulation even over sucrose rewards (Yoo et al. 2017). Beyond valence, cholinergic cells of the PTg have also been associated with arousal (Kobayashi et al. 2004; Kroeger et al. 2017) and attention (Inglis et al. 2001; Cyr et al. 2015), and both cholinergic and glutamatergic cells are associated with sleep-wake states (Kroeger et al. 2017). Consistent with this, lesions of cholinergic PTg neurons in similar tasks reduce accuracy and increase omissions, reflecting attentional deficits (Inglis et al. 2001). In our experiments, omission rates remained very low (<1%) across all stages, arguing against a generalized loss of task engagement. To test if transient attentional disruption or slowed cue processing could explain the performance deficit, we extended cue presentation to 3 s. Despite this, accuracy in stimulation trials remained at chance. Given that rodents can rapidly discriminate visual cues (200-320 ms for mice) (You (游文愷) and Mysore 2022), any slowing of perceptual processing would have to be substantial and sustained throughout the extended cue period to produce this effect. These results suggest that while brief attentional disruption may occur at stimulation onset, it is unlikely to account for the elimination of cue-guided performance. Instead, such pronounced behavioral effects may reflect combined recruitment of multiple neuronal populations in the PTg.

Consistent with the possibility that the behavioral phenotype reflects recruitment of multiple PTg populations, viral expression in our experiments was not restricted to a single neuronal population. Although the CamKIIa promoter has traditionally been considered to drive AAV expression primarily in excitatory neurons (Nathanson et al. 2009; Jones et al. 1994; Benson et al. 1992), its specificity varies across brain regions. In the rostral PTg, we observed substantial labelling of GABAergic neurons, along with smaller populations of glutamatergic and cholinergic neurons. This distribution aligns with previous studies of the cellular composition of the rostral PTg, with a high density of GABAergic cells, moderate density of glutamatergic cells and a sparse population of cholinergic cells (Luquin et al. 2018; Mena-Segovia et al. 2009). The PTg also contains subsets of neurons that co-express multiple neurotransmitters (Luquin et al. 2018; Mena-Segovia et al. 2009; Wang and Morales 2009). In our data, co-expression was observed between GAD2 and VGLUT2, whereas overlap between ChAT and GAD2 was minimal. Differences from previous reports of co-expression may reflect differential expression of GAD2 used in the present study and GAD1, which has been examined in prior work (Luquin et al. 2018; Wang and Morales 2009). Although co-expressing neurons were sparse and are unlikely to directly drive the behavioral effects, their presence highlights the cellular diversity of the PTg and suggests potential integration between excitatory and inhibitory signaling. GABAergic PTg neurons have also been implicated in disrupting goal-directed behavior through projections to the substantia nigra and basal ganglia (Gut et al. 2022; Hormigo et al. 2019; Hormigo, Zhou, Chabbert, et al. 2021), suggesting a possible substrate for the observed impairments. Together, this cellular and circuit-level heterogeneity supports a view of the PTg as more than a locomotor structure, with perturbations affecting both movement and the sensorimotor processes that guide action.

### Implications for sensorimotor gating and disease

The PTg has long been studied for its role in locomotion and as a potential therapeutic target in Parkinson’s disease (PD). Deep brain stimulation of the PTg has produced mixed results, with outcomes depending strongly on electrode placement (Hamani et al. 2016; Goetz et al. 2019). Though PD is classically associated with loss of dopaminergic neurons in the midbrain, particularly in the SNc, dopaminergic degeneration is considered part of the later stages of pathology, whereas loss of cholinergic cells in the PTg precedes the degeneration of cells in the SNc (Goetz et al. 2019; Vitale et al. 2019). Moreover, the PTg has been associated with several neurodegenerative diseases including Parkinson’s (PD), Alzheimer’s, supranuclear palsy (Vitale et al. 2019) and movement disorders, in particular through its association with freezing of gait (FOG) (Nutt et al. 2011).

FOG in PD patients has been shown to correlate with decreased connectivity between the PTg and other motor-related areas such as the thalamus, frontal cortex, and cerebellum, and to occur independently of dopaminergic degeneration (Chang et al. 2025). FOG is observed across multiple movement disorders and does not consistently correlate with core motor symptoms of PD such as tremor, bradykinesia, and rigidity (Nutt et al. 2011). Instead, FOG has been linked to impairments in executive function and perceptual processing, and its severity increases in cognitively demanding or visually complex environments (Nutt et al. 2011). For example, patients with FOG may struggle to appropriately adjust their gait when passing through a doorway, despite accurately judging the size of the opening when seated (Nutt et al. 2011). These findings suggest that FOG reflects a failure to appropriately integrate perceptual information with motor planning and execution, rather than a purely motor deficit (Nutt et al. 2011; Chang et al. 2025).

Psychotic symptoms can also occur in patients with PD, including hallucinations and delusions and cannot be fully explained by dopaminergic medication effects alone, as similar symptoms have been reported in drug-naïve patients (Pagonabarraga et al. 2024). Schizophrenia, a psychiatric disorder often associated with psychosis and executive dysfunction, has been linked to an increased risk of developing PD (Kuusimäki et al. 2021). Although PD and schizophrenia are often viewed as disorders at opposite ends on the spectrum of dopaminergic diseases, both involve impairments in perception and sensory processing. In particular, both conditions exhibit deficits in pre-pulse inhibition (Zhang et al. 2025; Dawson et al. 2000), a widely used measure of sensorimotor gating. Importantly, the PTg has been shown to contribute to pre-pulse inhibition (Diederich and Koch 2005), supporting the idea that dysfunction within this nucleus may contribute to overlapping motor, perceptual, and cognitive symptoms across neurological and psychiatric disease.

Together, these clinical observations align with our finding that rostral PTg stimulation does not merely suppress motor output but also disrupts the ability to use sensory cue information to guide action. This supports a broader view of the PTg as a sensorimotor interface, where disruption may affect not only movement execution but also the integration of perceptual information with ongoing and planned behavior.

### Limitations of the study

Because viral expression was not cell-type specific, we cannot attribute the observed effects to a particular neuronal population or projection. Optic fibers were positioned immediately dorsal to the rostral PTg and viral expression was concentrated in this region, but light spread and activation of transduced neurons near the anatomical border of the PTg cannot be fully excluded. This is particularly relevant for the SNr, which lies close to the stimulation site and has been implicated in ipsilateral orienting. We did not observe a consistent ipsilateral or contralateral response bias relative to the side of stimulation, arguing against a dominant contribution from direct recruitment of lateralized orienting circuits. Moreover, although SNr-related circuits have been implicated in gating signaled avoidance, this function is thought to be mediated through the PTg (Hormigo et al. 2019; Hormigo, Shanmugasundaram, Zhou, et al. 2021; Hormigo, Zhou, Chabbert, et al. 2021).

Beyond anatomical considerations, stimulation-induced motor arrest may also include suppression of eye movements, potentially altering visual sampling during cue presentation. Rats do not rely on foveation to the same extent as primates and can acquire visual information across a wide monocular field (Salinas-Navarro et al. 2009; Wallace et al. 2013; Zahler et al. 2023) but reduced orienting or visual scanning during stimulation remains a possible contributor. All sessions were video recorded and observed live, and although rare trials occurred in which rats turned away from the screen at stimulation onset, rats generally remained oriented toward the screen. These considerations limit our ability to distinguish between altered sensory acquisition and downstream cue-related processing. The precise disrupted step—cue detection, maintenance of cue information, action selection, or initiation of the selected response—therefore remains unresolved.

## Data and code availability

Data and code can be made available upon reasonable request.

## Acknowledgements

This work was funded by The Lundbeck Foundation (no. R366-2021-233, R.W.B) and the Novo Nordisk Foundation (no. NNF23OC0082192, R.W.B). The authors thank all members of the lab for their support.

Special thanks to Palle Koch Sørensen, who designed the Arduino - Raspberry Pi interface, our laser trigger software and for constructing the main frame of the behavioral chambers, and to Niels Thulin Johansen, who expanded the chamber hardware to include sound and light cues, and to Mikkel Roald-Arboel for his contribution in the early conceptual phase of the project.

## Author contributions

R.W.B., M.C.A.B., S.D.L, and R.J.F.S. conceptualization of the original idea. R.J.F.S. and M.C.A.B. developed the behavioral chamber hardware. S.D.L. developed the software. M.C.A.B. development of experimental protocol, rat training, data collection and surgeries.K.S. assisted with rat training, chamber maintenance and surgeries. K.S. and H.B. perfusion. G.H. 3D clearing and imaging. M.C.T. image analysis of cleared samples. O.D. RNAscope, imaging and image analysis. S.D.L. analysis of experimental data. M.C.A.B. writing original draft. S.D.L., M.C.T., K.S., and R.W.B. editing. R.W.B. supervision.

## Experimental model and study participant details

All animal experiments were approved by the Danish Animal Experiments inspectorate (2024-17315-0201-01739) and were in accordance with the EU Directive 2010/63/EU. 12 wild type male Long Evans rats (CRL:LE, 94-006-LE-M-06W) were pair housed in techniplast, top filtered, IVC cages (462 x 403 x 404 mm), in a room maintained at 22°C (± 2°C), 55% (± 10%) relative humidity and a 12-hour light/dark cycle with a 15–30-minute twilight period. The cages were lined with Aspen chip bedding (Tapvei, Vaikkojoentie 33, Kortteinen, Finland) and enriched with shredded paper as nesting material (Lilico PO Box 431, RH60UW, United Kingdom), a red plastic JAKO shelter (Molytex, Glostrup, Denmark), two Gnawing sticks (Tapvei, Vaikkojoentie 33, Kortteinen, Finland) and two cardboard tubes (Lilico PO Box 431, RH60UW, United Kingdom). For the first 7 days after arrival, all rats were acclimatized to the facility, with ad libitum access to food and water, but after acclimatization, the rats were habituated to handling, gradually food restricted and kept on a restricted diet (5 g per 100 g body weight) for training and data acquisition periods.

## Method details

### Stereotaxic surgery

12 hours before surgery, rats received a subcutaneous injection of extended-release Buprenorphine (0.65 mg/kg, Ethiqa XR 1.3 mg/ml), then Carprofen (0.01 ml/100 g, Rimadyl Bovis vet. 50 mg/ml) right before the surgery, and Lidocaine (10 mg/kg Lidocaine, 5 mg/kg Bupivacaine, Lidor vet. 20 mg/ml) before the first incision was made. Anesthesia was induced with 4% isoflurane (Attane vet. 1000 mg/g isoflurane) with an EZ anesthesia vaporizer in an induction chamber (EZ-178 sure-seal mouse/rat chamber), after which they received an intraperitoneal injection of premixed Ketamine/Xylazine (60 mg/kg Ketamine, 6 mg/kg Xylazine, Ketaminol Vet 20 mg/ml and Xysol Vet 2 mg/ml). The anesthesia was then maintained at 0.5-1.5% isoflurane. Anesthetic depth was regularly tested by checking withdrawal reflex of the hindlimb by toe pinch, and the rats’ HR, RR, temperature and perfusion were monitored with the PhysioSuite®. The surface of the skull was thoroughly cleaned, dried, and prepped with Kerr OptiBond FL, Primer and Adhesive, before mounting the base of a protective crown (PLA 3D print (Vöröslakos et al. 2021)), with UV curable dental cement (Tetric EvoFlow A1). To ensure flat skull position, Bregma and Lambda was brought to the same DV coordinate (<0.05 mm difference) as well as one point 2 mm lateral to the midline, on both sides. All rats were injected bilaterally in the rostral PTg (AP = -7.9 (+/-0.1), ML = 1.94 (+/-0.1), DV = 7.8 (+/- 0.1)) at 100 nl/min, with 500 nl of rAAV2/9-CamKIIa-ChrimsonR-mScarlet-KV2.1. pAAV-CamKIIa-ChrimsonR-mScarlet-KV2.1 was a gift from Christopher Harvey (Addgene viral prep # 124651-AAV9; http://n2t.net/addgene:124651; RRID:Addgene_124651)(Chettih and Harvey 2019). 10 minutes after the injection had finished, the capillary was slowly retracted at 1 mm/min and an optic fiber (RWD white ceramic ferrule Ø1.25, 200 μm core, 0.39 NA, length = 10 mm) was implanted 100 μm above each injection site. To secure the optic fibers, a thread was tied around the small indent of each ferrule. When both fibers had been secured onto the skull with UV curable dental cement (Tetric Evoflow A1), the ends of the strings were wrapped around the ferrules and embedded in a 2-component dental cement (Cenger, UNIFAST Trad powder and liquid). A scaffold for the crown was shaped using copper mesh and enforced by covering the surface with dental cement (Cenger, UNIFAST Trad powder and liquid).

To minimize post-operative inflammation, rats received a subcutaneous injection of Carprofen (0.01 ml/100g, Rimadyl Bovis vet. 50 mg/ml) every 24 hours for 3 days. Due to an increased risk of Pica behavior and constipation in rodents treated with Ethiqa XR, rats were housed in cages with soft tissue bedding and were fed 25 g of hydrating diet gel (clearH2O DietGel Recovery, Scanbur) and approximately 15 g of softened food pellets (dissolved in water) in addition to their usual dry food pellets, for the first 3 days after surgery. To minimize contact and damage to the implant and surrounding tissue, during postoperative recovery, rats were kept separate from their housing partner with a plastic cage divider, and the rat houses and cardboard tubes were removed for 2-4 days, depending on their rate of recovery, and instead had additional bedding and a soft nesting material (shredded tissue) during this period.

### Optogenetic stimulation

Optogenetic stimulation was delivered using a 625 nm laser (M625F2 Thorlabs) controlled by a pulse train generator (Serial Com interface for Arduino, by Palle Koch). Stimulation consisted of 10 ms pulses at 50 Hz, for 2.5–5.0 s depending on the stimulation stage. To avoid saturation of the neuronal response, and to have within subject controls, stimulation was delivered on 50% of the trials (per 10 trials) in a pseudorandomized order, ensuring stimulation did not occur more than three times in a row. Before behavioral testing, rats were screened for their response to optogenetic stimulation. The laser setting was adjusted to the lowest level that reliably evoked visible motor arrest across rats, defined as cessation of limb and head movements time-locked to light stimulation and observable by eye during live monitoring. This stimulation setting was then kept constant across behavioral testing. Light output was not measured at the patch cord or fiber tip before testing; therefore, absolute optical power cannot be reported.

### Behavioral Chambers

The chambers are 400x400x445 mm and made of polycarbonate, each with a bottom tray and a metal rod floor (**Supplementary Figure 1**). The chambers are equipped with a touchscreen (Keetouch 10.4”, KT-104-SP0S0M1) and an infrared (IR) curtain overlay (TopOneTech 10.4” IR touch screen, for response registration without screen touching), and the side opposite to the screen has a metal dispenser for delivery of a liquid reward, which drips into a small polycarbonate tub. Each chamber is equipped with a speaker, which delivers sound cues for correct (Sound Cue 1) and incorrect (Sound Cue 2) responses. A Plexiglas mask was placed on top of the screen and the IR overlay frame, to limit interaction with the screen to the relevant areas.

To enable automated training in the chambers, an interface was designed to bridge communication between an Arduino Mega2560 and a Raspberry Pi 4. The Raspberry Pi runs the VSCT program and is responsible for speaker, screen, IR curtain overlay, houselight, keyboard, and mouse functions, while the Arduino runs the reward delivery pump and triggers activation of the laser for optogenetic stimulation. The program is written in Python (Pygame module) and runs on the Raspberry Pi with the IDE, Thonny. Each chamber is individually controlled; variable parameters can be specified for the session and the resulting datafile can be named on the main screen of the program before starting a session. The chambers are placed inside a cabinet with 3 compartments (inner measurements of each compartment: h:672mm, w:558mm, d:578mm), each fitting one chamber, a USB camera (ELP-USB130W01MT-L21, iSPY software) for monitoring the rats live, and a pulley system which enables the rats to move to all areas of the chamber with minimal effort, without pulling on their fiber or building up strain in the patch cable (**Supplementary Figure 1**). It also ensures that the patch cable will be pulled out of reach of the rats. A commuter (RJ1 Thorlabs) is suspended from the ceiling of the chamber cabinets with a strong rubber band (Elastomer 18200-07 Fine science tools), and the bottom of the commuter was attached to the rats’ crown with crocodile clip on a string inside a thin rubber tube. Stimulation timing was coordinated by the VSCT program running on the Raspberry Pi. The Raspberry Pi controlled task-event timing and sent trigger commands at predefined points in the trial. These commands were relayed through the Arduino to a separate computer, which generated the output used to trigger the laser. Thus, stimulation timing was defined relative to the programmed trial events.

### VSCT Training Protocol

#### Food restriction and handling

Rats were gradually food restricted over three days from 10g per 100g body weight to 5g per 100g body weight. Food amounts were adjusted as the rats grew, and animals were weighed regularly to ensure that body weight did not fall below 85% of free-feeding body weight. Rats were weighed daily until 15 weeks of age, after which body weight was monitored weekly. During the first week of food restriction, rats were handled for at least 15 min daily per cage and fed drops of strawberry milk (Nestlé strawberry milk powder) with a syringe to habituate them to the trainer and to the reward used in the task.

#### Chamber Habituation & Conditioning

Each cage pair was assigned to a chamber at the beginning of the first habituation session, and rats were trained in the same order every day. This order was maintained throughout all training stages. Rats were habituated to the chambers and conditioned to Sound Cue 1, which signaled a reward (three drops of strawberry milk). At the start of each session, 10 drops of strawberry milk were available in the dispenser tub, and three additional drops were dispensed randomly within every 20 s. Sound Cue 1 was played simultaneously with each milk delivery to establish an association between the cue and the reward. A houselight was installed on top of each chamber to signal the premature response window. During habituation and Stages 1–2, the houselight remained on throughout each session. All rats completed three 20 min habituation sessions before progressing to Stage 1.

**Stage 1: Screen interaction.** In Stage 1, rats were taught to interact with the screen and Sound Cue 1 conditioning was further strengthened. A small white rectangle was displayed on the screen (**Supplementary Figure 2A**: Initiation button). When the rat approached closely enough to break the IR grid, Sound Cue 1 would trigger immediately, three drops of reward were dispensed, and the rectangle blinked three times. Rats were required to complete ≥80 trials in 30 minutes to progress to Stage 2.

**Stage 2: Trial initiation and stimulus-reward pairing**. In Stage 2 (**Supplementary Figure 2A**), the initiation button no longer elicited reward release but instead functioned as a trial initiation button. When the rat responded to the initiation button, the screen turned black for 1 s (black screen interval 1; BSI 1), after which a white square (visual stimulus) appeared in one of three positions on the screen. Responses to the visual stimulus, triggered a correct response and reward delivery. In this stage the visual stimulus was paired with the release of reward, and the rat learned that the initiation button no longer released reward, but was necessary to interact with, to get a visual stimulus to appear. To reduce side bias, the position of the visual stimulus was pseudorandomized, with no more than three consecutive presentations in the same position. If the rat did not respond within the 30 s limited hold period, an omission was recorded, and the trial was terminated. Rats were required to complete ≥80 trials in a 30 min session to progress to Stage 3.

**Stage 3: Cue introduction and response withholding.** In Stage 3 (**Supplementary Figure 2B**), rats were introduced to the visual cue, a small white plus sign, which appeared in one of the three positions. After trial initiation, there was a 0.5 s BSI 1, followed by a 0.5 s visual cue presentation. The cue appeared pseudorandomly across positions, as in Stage 2. Following the cue, there was a 0.5 s BSI 2, after which a visual stimulus was presented in the same position in which the cue had appeared. Touching the visual stimulus triggered a correct response. Because the goal was to teach rats that correct responses were tied to interaction with the visual stimulus, a premature-response window (PRW) was implemented during cue presentation. To signal the PRW, the houselight turned off during cue presentation. Interaction with the screen during the PRW would terminate the trial, trigger an incorrect response sound cue (sound cue 2) and a timeout. As rats learned to wait until after the cue disappeared, they also began to associate the position of the cue with the position of the subsequent stimulus. Rats were required to complete ≥40 correct trials in a session of 40 min to move on to Stage 4.

**Stage 4: Cue-guided choice.** In Stage 4 (**Supplementary Figure 2C**), rats learned to choose among three identical visual stimuli based on the position of the preceding cue. After trial initiation and cue presentation, three identical visual stimuli were presented, one in each position on the screen.

A correct response occurred when the rat selected the visual stimulus in the same position as the preceding visual cue while selection of either of the other two stimuli resulted in an incorrect response and a timeout. Cue position was pseudorandomized, but on this stage, incorrect trials triggered a correction trial, which repeated the same cue arrangement until the rat responded correctly. This procedure reduced the likelihood of rats adopting a fixed response strategy to obtain reward on approximately one third of the trials by chance. Rats were required to complete ≥40 correct trials in a 40 min session to progress to Stage 5.

**Stage 5: Final task shaping.** In Stage 5 correction trials were removed and several task parameters were gradually adjusted across three substages (**Supplementary Table 1**). In Stage 5.0, the intertrial interval increased from 3 s to 5 s and the limited hold decreased from 30 s to 20 s. In Stage 5.1 both black-screen intervals increased from 0.5 s to 1 s and the limited hold decreased to 10 s. Finally, in Stage 5.2 BSI 1 increased to 1.5 s and the limited hold decreased to 5 s. Rats were required to reach ≥60% accuracy with ≤20% premature responses with Stage 5.2 parameters before undergoing surgery.

#### Laser setup training

One week after the last rat of the cohort had undergone surgery, the rats were retrained on Stage 5.2 until reaching Stage 5.2 criteria again.

The rats were then trained on Stage 5.2 but with the laser attached to habituate them to the laser setup. Once a rat could reach ≥60% accuracy with ≤20% premature responses while attached to the laser setup, it progressed to the stimulation stages (Stages 6-9).

#### VSCT task parameters and outcome definitions

Correct and incorrect trials were used to calculate session accuracy and served as the primary measure of task performance.

A *correct trial* was recorded when the rat chose the visual stimulus in the same position as the preceding cue. This triggered a correct response, which activated Sound Cue 1, caused the chosen visual stimulus to blink 3 times, and initiated reward delivery.

An *incorrect trial* was recorded when the rat chose one of the two visual stimuli that did not match the cue position. Incorrect response resulted in immediate trial termination, activation of Sound Cue 2, and a timeout.

A *timeout* consisted of a 5 s period during which the screen turned red and the rat was unable to initiate a new trial. Timeouts occurred following incorrect responses or premature responses.

*Premature responses (PR)* were responses that occurred during the *premature response window (PRW)*, defined as the period within a trial when interaction with the screen was not allowed. PRs were excluded from accuracy calculations but were used as a measure of impulsivity and task related frustration. Like incorrect trials, PRs terminated the trial, triggered Sound Cue 2 and resulted in a timeout.

The *intertrial interval (ITI)* was the time between the end of a trial and the beginning of the next.

The *limited hold (LH)* was the time window during which the rat was required to respond to the stimuli. Failure to respond within the LH resulted in an omission, after which the next trial began following the ITI. Omissions were used as a measure of task engagement or motivation and were excluded from accuracy calculations.

### Data collection

Two full laser setups were assembled so stimulation sessions could run in two chambers simultaneously. Rats were recorded from in the same order every day, but on some days the laser would be attached to their left optic fiber and on other days their right.

The laser setup can be distracting and stressful for some rats, thus not all rats in the cohort were able to reach 60% accuracy and/or have 50% premature responses and were therefore not included in the final analysis. All rats went through 1-2 sessions a day of approximately 30 minutes per stage. Animals progressed through stimulation stages according to their behavioral performance and welfare. Stage order was adjusted when necessary to minimize frustration and maintain task engagement.

Even though all sessions were video recorded the rats were closely monitored at all times to make sure any incidents could be responded to immediately and to track the performance of the rats live. Rats that did well on their first session were moved to a different stage after, for a consecutive session of a different stage until all rats had completed 4 sessions on each stage with ≥50 response trials (Om and PR excluded).

### Quantification and statistical analysis

A session was included for quantification if the accuracy in control trials was above 60% and it included at least 50 trials that were not a premature response or an omission. Accuracy was defined as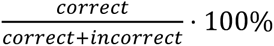 and mean side bias was defined as:

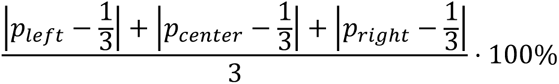

with p being the proportion of responses at a given location.

When simulating randomness for comparison to real sessions, a session with the same total number of trials was created and each response in the list was assigned either correct or incorrect independently and randomly with a probability of 0.666 and 0.334 respectively.

All summary quantifications are given by mean ± standard error of the mean (SEM) unless stated otherwise. All statistical tests are non-parametric as the sample sizes were not determined to be large enough for quantifications of normality or assumptions of variance distributions. As all statistical quantifications were between different states of the same cohort of rats, a paired design was employed, using a two-sided Wilcoxon signed-rank test for single comparisons. For comparisons of more than two groups the Friedman test was used. The null hypothesis was rejected if p<0.05. Cohort size was chosen to ensure that at least 6 rats would be successfully implanted and survive until quantification as this the minimum sample size required for p<0.05 in a Wilcoxon signed-rank test. This was considered a compromise between minimizing animal use and assuring statistical power. No randomization or blinding was considered necessary for an unbiased design given the design’s internal control and the automated nature of data acquisition.

### Dissection and fixation

All rats were sacrificed by transcardiac perfusion with 35-40 ml Phosphate-buffered saline (PBS) followed by 40-45 ml 4% Paraformaldehyde (PFA). The perfused brain tissues were placed in separate 50 ml Falcon tubes and post-fixed for 24 hours in 15-20 ml 4% PFA. After post-fixation, six of the brains were cut along the midline and placed in separate Falcon tubes, with one hemisphere used for 3D clearing, and the other for RNAscope.

#### Whole-mount immunolabelling and optical clearing of rat central nervous system tissue

Rat brains were processed using a modified Adipo-Clear+ (Branch et al. 2021) whole-mount immunolabelling and solvent-based optical clearing protocol. Following post-fixation, samples were subsequently washed several times in phosphate-buffered saline (PBS) containing 0.02% sodium azide.

To reduce tissue pigmentation and improve antibody penetration, samples were incubated in 25% Quadrol in water for 2 days at 37°C, followed by 5% ammonium hydroxide in water for 1 day at 37°C. Samples were then washed in B1n buffer for 1 day at room temperature. B1n buffer consisted of 0.1% Triton X-100, 4% glycine, 0.001% sodium azide and 1 mM NaOH in water.

Samples were dehydrated at room temperature through a graded methanol series consisting of 20%, 40% and 60% methanol in B1n buffer, followed by 80% methanol in water. Each step was performed for 1.5 h. Samples were subsequently incubated twice in 100% methanol for 1 h per incubation, followed by overnight incubation in dichloromethane: methanol at a 2:1 ratio. After several washes in 100% methanol, endogenous fluorescence and residual pigments were bleached by incubation in 5% hydrogen peroxide in methanol for 2 days at 4°C.

Samples were rehydrated at room temperature through 80%, 60%, 40% and 20% methanol in water for 1.5 h per step and were subsequently washed in PBS for 1 h. Tissues were then incubated in SdC solution for 1 day at 37°C. SdC solution contained 1% 2-hydroxypropyl-β-cyclodextrin (PanReac AppliChem, A0367,0100), 1× PBS and 0.02% sodium azide in water. Samples were permeabilized overnight, at 37°C in PTwH (1x PBS 0.2% tween20 and 10 µg/mL heparin) supplemented with 5% dimethyl sulfoxide and 0.3 M glycine.

For immunolabelling, tissues were blocked for 1 day at 37°C in PTxH (1x PBS 0.2% triton and 10 µg/mL heparin) containing 3% normal donkey serum. Samples were incubated with primary antibodies (diluted in blocking solution for 14–28 days at 37°C). Primary antibodies included antibodies against choline acetyltransferase (ChAT, CHEMICON, item no. AB144P, concentration 1:400), red fluorescent protein (RFP, Rockland, Item No.600-401-379, concentration 1:1000) and green fluorescent protein (GFP, Aves Labs, SKU: GFP-1020, concentration 1:3000), depending on the fluorescent reporter expressed in each animal. Following primary antibody incubation, samples were washed extensively in PTwH for at least 2 days at 37°C. PTwH. Samples were then incubated for 14 days at 37°C with species-appropriate donkey secondary antibodies (Alexa Fluor 647, 555, and 488) from Jackson Immuno Research diluted 1:1000 in PTwH. Fluorophore combinations were selected according to the reporter proteins present in each sample. RFP and GFP immunoreactivity were generally detected using far-red fluorophore-conjugated secondary antibodies (Alexa Fluor 647), whereas ChAT was detected using a fluorophore emitting in the orange-red range (Alexa Fluor 555). For samples requiring alternative channel separation, GFP and ChAT were detected using orange-red and green fluorophores (Alexa Fluor 488), respectively. After secondary antibody incubation, samples were washed in PTwH for at least 2 days, typically over a weekend. Samples were then dehydrated at room temperature in 20%, 40% and 60% methanol in PBS for 1.5 h per step, followed by 80% methanol in PBS for 2 h. Tissues were incubated in 100% methanol for 2 h and then overnight in fresh 100% methanol.

For lipid extraction and refractive-index matching, samples were incubated in dichloromethane:methanol at a 2:1 ratio, first for one wash and subsequently overnight at room temperature. Samples were then transferred to 100% dichloromethane for two consecutive 2-h incubations followed by an overnight incubation. Finally, tissues were transferred to ethyl cinnamate and stored in ethyl cinnamate at room temperature until imaging. The ethyl cinnamate was replaced at least once after initial clearing. All incubations were performed with gentle agitation unless otherwise stated.

Imaging was done using an Ultramicroscope Blaze (Miltenyi Biotec) lightsheet microscope. The microscope was installed on an air table to reduce vibration artifacts. There are two illumination objectives (0.0135–0.135 NA) that generate 3 bidirectional light sheets together with a 4x and 12x detection objective (MIPLAN 4x: 0.35 NA, WD 16 mm and MI PLAN 12x: 0.53 NA, WD 10.9 mm) without zoom. The images were acquired with a 5.5 Mpx sCMOS camera, pixel size 6.5 μ m using ImSpector software, in fast tiling “lightspeed mode”. The laser line used to capture either the 488, 555, or 647 signal (which was enhanced with an Alexa Fluor 647, 555, or 488 secondary antibody) was a 488, 561, or 639 nm, diode laser.

### RNAscope

To investigate the viral distribution in the main neuronal populations of the PTg, we used multiplex RNAscope on sagittal sections of the PTg and surrounding area, targeting VGLUT2 (vesicular glutamate transporter 2), ChAT (choline acetyl transferase), and GAD2 (glutamate decarboxylase 2). The analysis was based on one hemisphere from six different animals and the area of the PTg was determined by its cholinergic neurons in each section. Only cells expressing the virus within this area (14189 cells) were included in the analysis.

For cryoprotection, tissues were incubated for up to three days in PBS with 30% sucrose (until they sank to the bottom of the tube), then frozen and stored at -80°C for approximately one month. The whole hemisphere was used for cryosectioning, as more structures can be used to recognize the area of the PTg. Fixed-frozen rat brain sections (12µm sections on Superfrost™ slides) were processed for RNAscope labelling followed by immunohistochemistry (IHC). Glutamatergic, cholinergic and GABAergic neurons were identified using the RNAscope Multiplex Fluorescent Assay V2 (Advanced Cell Diagnostics) with the probes: Slc17a6-C2 (317011-C2), Chat-C1 (430111), and Gad2-C3 (316401-C3), according to the manufacturer’s instructions. Tissue quality was assessed using the RNAscope Positive 3-plex control probe (320891) and Negative 3-plex control probe (320871). To detect neurons that expressed the RFP plasmid, sections were blocked with 5% donkey serum following RNAscope labelling and incubated overnight with an anti-RFP antibody (1:500, Rockland). After washing sections were incubated with Alexa Fluor 568-conjugated Donkey Anti-Mouse IgG (Invitrogen) and mounted using ProLong™ Diamond Antifade Mountant (Invitrogen). Imaging was performed on a Carl Zeiss AxioImager microscope through a Plan-Apochromat 20x/0.8 objective. The PTg was identified on sagittal sections by ChAT labeling: ChAT-positive neurons form a characteristic triangular cluster on sagittal planes, and the border of this triangle was used to define the PTg ROI. QuPath was used to delineate ROI based on ChAT staining. Within this ROI all RFP-positive neurons were detected and segmented, and the mean fluorescent intensity per cell was measured for ChAT, GAD2 and Slc17a6. A fluorescent intensity value higher than 1000 AU was used as a threshold to determine if a cell expressed the corresponding neurotransmitter marker. To reduce the number of false positives in the co-expression analysis the threshold was increased to 2000 AU.

## References

Azzopardi, Erin, Andrea G. Louttit, Cleusa DeOliveira, Steven R. Laviolette, and Susanne Schmid. 2018. “The Role of Cholinergic Midbrain Neurons in Startle and Prepulse Inhibition.” The Journal of Neuroscience 38 (41): 8798–808. 10.1523/JNEUROSCI.0984-18.2018.

Benson, D. L., P. J. Isackson, C. M. Gall, and E. G. Jones. 1992. “Contrasting Patterns in the Localization of Glutamic Acid Decarboxylase and Ca2+/Calmodulin Protein Kinase Gene Expression in the Rat Central Nervous System.” Neuroscience 46: 825–49. DOI:%2010.1016/0306-4522(92)90188-8.

Branch, Audrey, Daniel Tward, Anna C. Kolstad, et al. 2021. “An Optimized Tissue Clearing Protocol for Rat Brain Labeling, Imaging, and High Throughput Analysis.” Preprint, bioRxiv, March 27. 10.1101/639674.

Caggiano, V., R. Leiras, H. Goñi-Erro, et al. 2018. “Midbrain Circuits That Set Locomotor Speed and Gait Selection.” Nature 553 (7689): 455–60. 10.1038/nature25448.

Carvalho, Miguel M., Nouk Tanke, Emilio Kropff, Menno P. Witter, May-Britt Moser, and Edvard I. Moser. 2020. “A Brainstem Locomotor Circuit Drives the Activity of Speed Cells in the Medial Entorhinal Cortex.” Cell Reports 32 (10). 10.1016/j.celrep.2020.108123.

Chang, Hee Jin, Huijin Song, Chul-Ho Sohn, and Han-Joon Kim. 2025. “Altered Pedunculopontine Pathways in Freezing of Gait: A Diffusion Tensor Imaging Study Based on Dopaminergic Degeneration Status.” Journal of Neurology 272 (12): 777. 10.1007/s00415-025-13521-2.

Chettih, Selmaan N., and Christopher D. Harvey. 2019. “Single-Neuron Perturbations Reveal Feature-Specific Competition in V1.” Nature 567 (7748): 334–40. 10.1038/s41586-019-0997-6.

Cyr, Marilyn, Maxime J. Parent, Naguib Mechawar, et al. 2015. “Deficit in Sustained Attention Following Selective Cholinergic Lesion of the Pedunculopontine Tegmental Nucleus in Rat, as Measured with Both Post-Mortem Immunocytochemistry and in Vivo PET Imaging with [18F]Fluoroethoxybenzovesamicol.” Behavioural Brain Research 278 (February): 107–14. 10.1016/j.bbr.2014.09.021.

Dawson, Michael E., Anne M. Schell, Erin A. Hazlett, Keith H. Nuechterlein, and Diane L. Filion. 2000. “On the Clinical and Cognitive Meaning of Impaired Sensorimotor Gating in Schizophrenia.” Psychiatry Research 96 (3): 187–97. 10.1016/S0165-1781(00)00208-0.

Diederich, Kai, and Michael Koch. 2005. “Role of the Pedunculopontine Tegmental Nucleus in Sensorimotor Gating and Reward-Related Behavior in Rats.” Psychopharmacology 179 (2): 402–8. 10.1007/s00213-004-2052-y.

Goetz, Laurent, Manik Bhattacharjee, Murielle U. Ferraye, et al. 2019. “Deep Brain Stimulation of the Pedunculopontine Nucleus Area in Parkinson Disease: MRI-Based Anatomoclinical Correlations and Optimal Target.” Neurosurgery 84 (2): 506–18. 10.1093/neuros/nyy151.

Goñi-Erro, Haizea, Raghavendra Selvan, Vittorio Caggiano, Roberto Leiras, and Ole Kiehn. 2023. “Pedunculopontine Chx10+ Neurons Control Global Motor Arrest in Mice.” Nature Neuroscience 26 (9): 1516–28. 10.1038/s41593-023-01396-3.

Gut, Nadine K., Duygu Yilmaz, Krishnakanth Kondabolu, Icnelia Huerta-Ocampo, and Juan Mena-Segovia. 2022. “Selective Inhibition of Goal-Directed Actions in the Mesencephalic Locomotor Region.” Preprint, bioRxiv, January 20. 10.1101/2022.01.18.476772.

Hamani, Clement, Tipu Aziz, Bastiaan R. Bloem, et al. 2016. “Pedunculopontine Nucleus Region Deep Brain Stimulation in Parkinson Disease: Surgical Anatomy and Terminology.” Stereotactic and Functional Neurosurgery 94 (5): 298–306. 10.1159/000449010.

Hormigo, Sebastian, Bharanidharan Shanmugasundaram, Ji Zhou, and Manuel A. Castro-Alamancos. 2021. “A Signaled Locomotor Avoidance Action Is Fully Represented in the Neural Activity of the Midbrain Tegmentum.” The Journal of Neuroscience 41 (19): 4262–75. 10.1523/JNEUROSCI.0027-21.2021.

Hormigo, Sebastian, German Vega-Flores, Victor Rovira, and Manuel A. Castro-Alamancos. 2019. “Circuits That Mediate Expression of Signaled Active Avoidance Converge in the Pedunculopontine Tegmentum.” Research Articles. Journal of Neuroscience 39 (23): 4576–94. 10.1523/JNEUROSCI.0049-19.2019.

Hormigo, Sebastian, Ji Zhou, and Manuel A. Castro-Alamancos. 2021. “Bidirectional Control of Orienting Behavior by the Substantia Nigra Pars Reticulata: Distinct Significance of Head and Whisker Movements.” Research Article: New Research. eNeuro 8 (5). 10.1523/ENEURO.0165-21.2021.

Hormigo, Sebastian, Ji Zhou, Dorian Chabbert, Bharanidharan Shanmugasundaram, and Manuel A. Castro-Alamancos. 2021. “Basal Ganglia Output Has a Permissive Non-Driving Role in a Signaled Locomotor Action Mediated by the Midbrain.” Research Articles. Journal of Neuroscience 41 (7): 1529–52. 10.1523/JNEUROSCI.1067-20.2020.

Huang, Yanwang, Shangyi Wang, Qingxiu Wang, et al. 2024. “Glutamatergic Circuits in the Pedunculopontine Nucleus Modulate Multiple Motor Functions.” Neuroscience Bulletin 40 (11): 1713–31. 10.1007/s12264-024-01314-y.

Inagaki, Hidehiko K., Susu Chen, Margreet C. Ridder, et al. 2022. “A Midbrain-Thalamus-Cortex Circuit Reorganizes Cortical Dynamics to Initiate Movement.” Cell 185 (6): 1065–1081.e23. 10.1016/j.cell.2022.02.006.

Inglis, Wendy L., Mary C. Olmstead, and Trevor W. Robbins. 2001. “Selective Deficits in Attentional Performance on the 5-Choice Serial Reaction Time Task Following Pedunculopontine Tegmental Nucleus Lesions.” Behavioural Brain Research 123 (2): 117–31. 10.1016/S0166-4328(01)00181-4.

Jones, Eg, Gw Huntley, and Dl Benson. 1994. “Alpha Calcium/Calmodulin-Dependent Protein Kinase II Selectively Expressed in a Subpopulation of Excitatory Neurons in Monkey Sensory- Motor Cortex: Comparison with GAD-67 Expression.” The Journal of Neuroscience 14 (2): 611–29. 10.1523/JNEUROSCI.14-02-00611.1994.

Kaur, Jaspreet, Salif A. Komi, Oksana Dmytriyeva, Grace A. Houser, Madelaine C. A. Bonfils, and Rune W. Berg. 2025. “Pedunculopontine-Stimulation Obstructs Hippocampal Theta Rhythm and Halts Movement.” Scientific Reports 15 (1): 17903. 10.1038/s41598-025-01695-8.

Kobayashi, T., C. Good, K. Mamiya, R. D. Skinner, and E. Garcia-Rill. 2004. “Development of REM Sleep Drive and Clinical Implications.” Journal of Applied Physiology (Bethesda, Md. : 1985) 96 (2): 735–46. 10.1152/japplphysiol.00908.2003.

Kozlowska, Kasia, Peter Walker, Loyola McLean, and Pascal Carrive. 2015. “Fear and the Defense Cascade: Clinical Implications and Management.” Harvard Review of Psychiatry 23 (4): 263–87. 10.1097/HRP.0000000000000065.

Kroeger, Daniel, Loris L. Ferrari, Gaetan Petit, et al. 2017. “Cholinergic, Glutamatergic, and GABAergic Neurons of the Pedunculopontine Tegmental Nucleus Have Distinct Effects on Sleep/Wake Behavior in Mice.” The Journal of Neuroscience 37 (5): 1352–66. 10.1523/JNEUROSCI.1405-16.2016.

Kroeger, Daniel, Jack Thundercliffe, Alex Phung, et al. 2022. “Glutamatergic Pedunculopontine Tegmental Neurons Control Wakefulness and Locomotion via Distinct Axonal Projections.” Sleep 45 (12): zsac242. 10.1093/sleep/zsac242.

Kuusimäki, Tomi, Haidar Al-Abdulrasul, Samu Kurki, et al. 2021. “Increased Risk of Parkinson’s Disease in Patients With Schizophrenia Spectrum Disorders.” Movement Disorders 36 (6): 1353–61. 10.1002/mds.28484.

Lee, A. Moses, Jennifer L. Hoy, Antonello Bonci, Linda Wilbrecht, Michael P. Stryker, and Cristopher M. Niell. 2014. “Identification of a Brainstem Circuit Regulating Visual Cortical State in Parallel with Locomotion.” Neuron 83 (2): 455–66. 10.1016/j.neuron.2014.06.031.

Luquin, Esther, Ibone Huerta, María S. Aymerich, and Elisa Mengual. 2018. “Stereological Estimates of Glutamatergic, GABAergic, and Cholinergic Neurons in the Pedunculopontine and Laterodorsal Tegmental Nuclei in the Rat.” Frontiers in Neuroanatomy 12 (May). 10.3389/fnana.2018.00034.

Masini, Débora, and Ole Kiehn. 2022. “Targeted Activation of Midbrain Neurons Restores Locomotor Function in Mouse Models of Parkinsonism.” Nature Communications 13 (1): 504. 10.1038/s41467-022-28075-4.

Mena-Segovia, J., B. r. Micklem, R. g. Nair-Roberts, M. a. Ungless, and J. p. Bolam. 2009. “GABAergic Neuron Distribution in the Pedunculopontine Nucleus Defines Functional Subterritories.” Journal of Comparative Neurology 515 (4): 397–408. 10.1002/cne.22065.

Mosher, Clayton P., Adam N. Mamelak, Mahsa Malekmohammadi, Nader Pouratian, and Ueli Rutishauser. 2021. “Distinct Roles of Dorsal and Ventral Subthalamic Neurons in Action Selection and Cancellation.” Neuron 109 (5): 869–881.e6. 10.1016/j.neuron.2020.12.025.

Muthusamy, Kalai A., Bhooma R. Aravamuthan, Morten L. Kringelbach, et al. 2007. “Connectivity of the Human Pedunculopontine Nucleus Region and Diffusion Tensor Imaging in Surgical Targeting.” Journal of Neurosurgery. Journal of Neurosurgery 107 (4): 814–20. 10.3171/JNS-07/10/0814.

Nathanson, Jason L., Yuchio Yanagawa, Kunihiko Obata, and Edward M. Callaway. 2009. “Preferential Labeling of Inhibitory and Excitatory Cortical Neurons by Endogenous Tropism of AAV and Lentiviral Vectors.” Neuroscience 161 (2): 441– 50. 10.1016/j.neuroscience.2009.03.032.

Noftz, William A., Nichole L. Beebe, Jeffrey G. Mellott, and Brett R. Schofield. 2020. “Cholinergic Projections From the Pedunculopontine Tegmental Nucleus Contact Excitatory and Inhibitory Neurons in the Inferior Colliculus.” Frontiers in Neural Circuits 14 (July). 10.3389/fncir.2020.00043.

Nutt, John G., Bastiaan R. Bloem, Nir Giladi, Mark Hallett, Fay B. Horak, and Alice Nieuwboer. 2011. “Freezing of Gait: Moving Forward on a Mysterious Clinical Phenomenon.” The Lancet. Neurology 10 (8): 734–44. 10.1016/S1474-4422(11)70143-0.

Olszewski, J., and D. Baxter. 1982. Cytoarchitecture of the Human Brain Stem. Karger. https://books.google.dk/books?id=2GJnQgAACAAJ.

Özkan, Mazhar, Büşra Köse, Oktay Algın, Sinem Oğuz, Mert Emre Erden, and Safiye Çavdar. 2022. “Non-Motor Connections of the Pedunculopontine Nucleus of the Rat and Human Brain.” Neuroscience Letters 767 (January): 136308. 10.1016/j.neulet.2021.136308.

Pagonabarraga, Javier, Helena Bejr-Kasem, Saul Martinez-Horta, and Jaime Kulisevsky. 2024. “Parkinson Disease Psychosis: From Phenomenology to Neurobiological Mechanisms.” Nature Reviews Neurology 20 (3): 135–50. 10.1038/s41582-023-00918-8.

Pan, Wei-Xing, and Brian I. Hyland. 2005. “Pedunculopontine Tegmental Nucleus Controls Conditioned Responses of Midbrain Dopamine Neurons in Behaving Rats.” Behavioral/Systems/Cognitive. Journal of Neuroscience 25 (19): 4725–32. 10.1523/JNEUROSCI.0277-05.2005.

Roseberry, Thomas K., A. Moses Lee, Arnaud L. Lalive, Linda Wilbrecht, Antonello Bonci, and Anatol C. Kreitzer. 2016. “Cell-Type-Specific Control of Brainstem Locomotor Circuits by Basal Ganglia.” Cell 164 (3): 526–37. 10.1016/j.cell.2015.12.037.

Rye, David B., Clifford B. Saper, Henry J. Lee, and Bruce H. Wainer. 1987. “Pedunculopontine Tegmental Nucleus of the Rat: Cytoarchitecture, Cytochemistry, and Some Extrapyramidal Connections of the Mesopontine Tegmentum.” Journal of Comparative Neurology 259 (4): 483–528. 10.1002/cne.902590403.

Salinas-Navarro, M., S. Mayor-Torroglosa, M. Jiménez-López, et al. 2009. “A Computerized Analysis of the Entire Retinal Ganglion Cell Population and Its Spatial Distribution in Adult Rats.” Vision Research 49 (1): 115–26. 10.1016/j.visres.2008.09.029.

Schmidt, Robert, Daniel K. Leventhal, Nicolas Mallet, Fujun Chen, and Joshua D. Berke. 2013. “Canceling Actions Involves a Race between Basal Ganglia Pathways.” Nature Neuroscience 16 (8): 1118–24. 10.1038/nn.3456.

Shik, M. L., F. V. Severin, and G. N. Orlovskii. 1966. “Control of Walking and Running by Means of Electrical Stimulation of the Mid-Brain.” Biophysics (Russian Federation) 11 (4): 756–65.

Skinner, R. D., and E. Garcia-Rill. 1984. “The Mesencephalic Locomotor Region (MLR) in the Rat.” Brain Research 323 (2): 385–89. 10.1016/0006-8993(84)90319-6.

Tello, Andrea Juárez, Cornelis Immanuel van der Zouwen, Léonie Dejas, et al. 2024. “Dopamine-Sensitive Neurons in the Mesencephalic Locomotor Region Control Locomotion Initiation, Stop, and Turns.” Cell Reports 43 (5). 10.1016/j.celrep.2024.114187.

Thompson, John A., and Gidon Felsen. 2013. “Activity in Mouse Pedunculopontine Tegmental Nucleus Reflects Action and Outcome in a Decision-Making Task.” Journal of Neurophysiology 110 (12): 2817–29. 10.1152/jn.00464.2013.

Vitale, F., A. Capozzo, P. Mazzone, and E. Scarnati. 2019. “Neurophysiology of the Pedunculopontine Tegmental Nucleus.” Neurobiology of Disease 128 (August): 19–30. 10.1016/j.nbd.2018.03.004.

Vöröslakos, Mihály, Hiroyuki Miyawaki, Sebastien Royer, et al. 2021. “3D-Printed Recoverable Microdrive and Base Plate System for Rodent Electrophysiology.” Bio-Protocol 11 (16). 10.21769/BioProtoc.4137.

Wallace, Damian J., David S. Greenberg, Juergen Sawinski, Stefanie Rulla, Giuseppe Notaro, and Jason N. D. Kerr. 2013. “Rats Maintain an Overhead Binocular Field at the Expense of Constant Fusion.” Nature 498 (7452): 65–69. 10.1038/nature12153.

Wang, Hui-Ling, and Marisela Morales. 2009. “Pedunculopontine and Laterodorsal Tegmental Nuclei Contain Distinct Populations of Cholinergic, Glutamatergic and GABAergic Neurons in the Rat.” The European Journal of Neuroscience 29 (2): 10.1111/j.1460-9568.2008.06576.x. 10.1111/j.1460-9568.2008.06576.x.

Watson, Charles, Caitlin Bartholomaeus, and Luis Puelles. 2019. “Time for Radical Changes in Brain Stem Nomenclature—Applying the Lessons From Developmental Gene Patterns.” Frontiers in Neuroanatomy 13 (February). 10.3389/fnana.2019.00010.

Wolf, Andrew B., Mario J. Lintz, Jamie D. Costabile, John A. Thompson, Elizabeth A. Stubblefield, and Gidon Felsen. 2015. “An Integrative Role for the Superior Colliculus in Selecting Targets for Movements.” Journal of Neurophysiology 114 (4): 2118–31. 10.1152/jn.00262.2015.

Xiao, Cheng, Jounhong Ryan Cho, Chunyi Zhou, et al. 2016. “Cholinergic Mesopontine Signals Govern Locomotion and Reward through Dissociable Midbrain Pathways.” Neuron 90 (2): 333–47. 10.1016/j.neuron.2016.03.028.

Yau, Hau-Jie, Dong V. Wang, Jen-Hui Tsou, et al. 2016. “Pontomesencephalic Tegmental Afferents to VTA Non-Dopamine Neurons Are Necessary for Appetitive Pavlovian Learning.” Cell Reports 16 (10): 2699–710. 10.1016/j.celrep.2016.08.007.

Yoo, Ji Hoon, Vivien Zell, Johnathan Wu, et al. 2017. “Activation of Pedunculopontine Glutamate Neurons Is Reinforcing.” Research Articles. Journal of Neuroscience 37 (1): 38–46. 10.1523/JNEUROSCI.3082-16.2016.

You (游文愷), Wen-Kai, and Shreesh P. Mysore. 2022. “Dynamics of Visual Perceptual Decision-Making in Freely Behaving Mice.” eNeuro 9 (2): ENEURO.0161-21.2022. 10.1523/ENEURO.0161-21.2022.

Zahler, Sebastian H., David E. Taylor, Brennan S. Wright, et al. 2023. “Hindbrain Modules Differentially Transform Activity of Single Collicular Neurons to Coordinate Movements.” Cell 186 (14): 3062–3078.e20. 10.1016/j.cell.2023.05.031.

Zhang, Zhen, Ling Zhang, Xiaofeng Huang, et al. 2025. “Elucidation of Blink Reflex Characteristics in Parkinson’s Disease Subtypes through Prepulse Inhibition.” Frontiers in Neurology 16 (May). 10.3389/fneur.2025.1505598.

