## Supplementary material for "Cue-guided performance is disrupted during pedunculotegmental-induced motor arrest": Supplementary material.pdf

### Supplementary figure 1

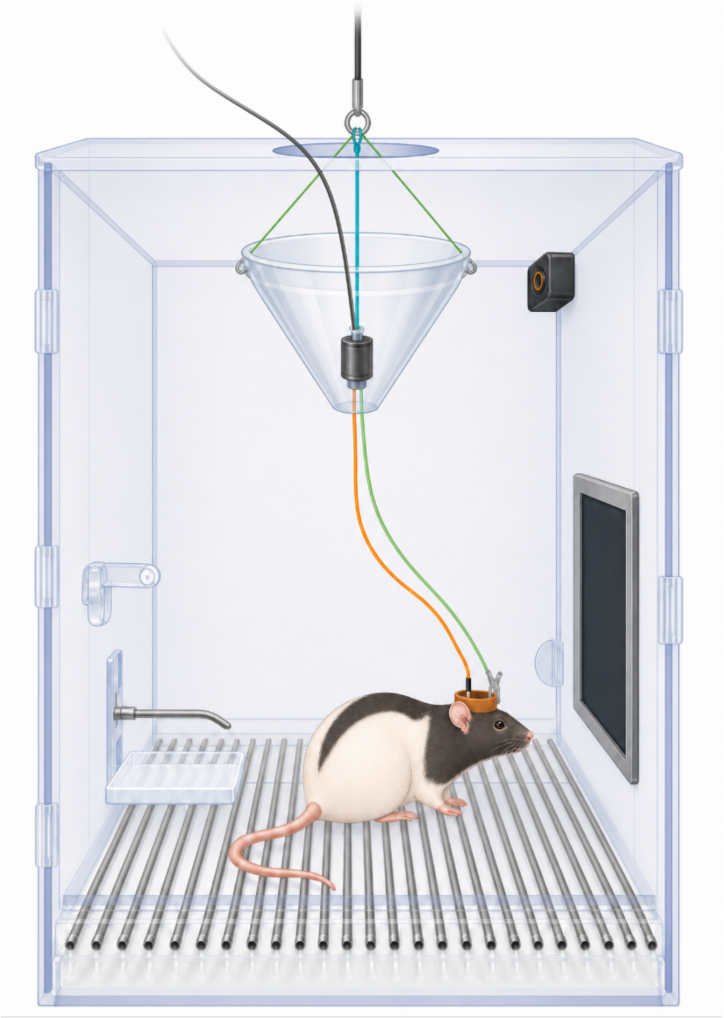

***Supplementary figure 1.*** *Chamber and laser setup. Left side: reward delivery polycarbonate tub and metal dispenser. Right: Touchscreen. Upper right corner: speaker. Laser setup: rat with chronically implanted optic fiber and a protective dental cement crown (brown). A crocodile clip is attached to the front of the crown and to the bottom of the rotary joint (black) via a string (green) to reduce strain on the patch cable (orange). Another patch cable (black) is attached to the top of the rotary joint and is plugged into a LED laser outside of the chambers. The rotary joint is suspended from the cabinet ceiling to allow the rat to move to all parts of the chamber and to pull away the patch cable when the rat is rearing. To ensure the rat cannot jump out of the chamber and that the rotary joint won't get caught on the edge of the hole in the chamber ceiling, a plastic cone was suspended from the chamber ceiling around the rotary joint.*

### Supplementary figure 2

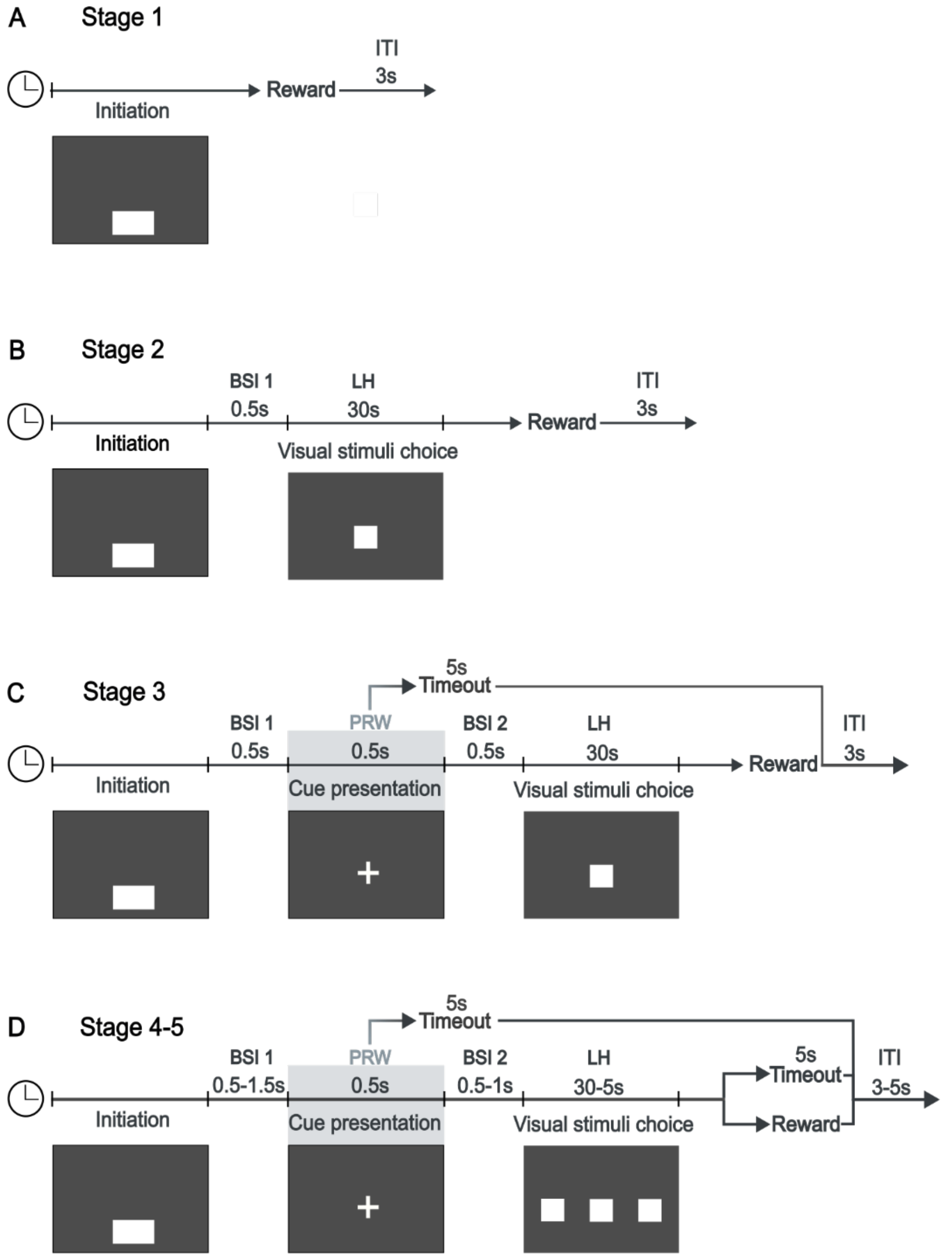


*Supplementary figure 2. Training stage overview. A) Stage 1, the rats learn to interact with the screen. Interaction with the initiation button elicits a reward, the button blinks, and sound cue 1 is played. B) Stage 2, the initiation button is followed by a black screen interval (BSI 1), then, a single visual stimulus, which will pseudorandomly appear in one of three positions on the screen. Only, interaction with the visual stimulus elicits a reward, blinking stimulus and sound cue 1. C) Stage 3, the cue is introduced. After initiation and BSI 1 a cue appears pseudorandomly in one of the three positions on the screen. During the presentation of the cue, the rats are not allowed to interact with the screen. Any touches registered during this time are recorded as premature responses and will trigger sound cue 2 (buzzer sound) and a timeout. After the cue, there is another black screen interval (BSI 2), followed by a visual stimulus in the same position as the preceding cue. Only responses to the visual stimulus will elicit a reward, blinking stimulus and sound cue 1. D) Stage 4-5, the rats must choose the correct visual stimulus. In both Stage 4 and 5 the cue is followed by three visually identical stimuli, but only the stimulus in the same position as the preceding cue is correct. In Stage 4, incorrect responses will lead to a "correction trial" where the cue will be in the same position as the previous trial. Stage 5 is subdivided into 5.0, 5.1, and 5.2 where the BSI 1, BSI 2, LH and ITI are gradually changed to the final version. ITI = Inter trial interval, BSI = Black screen interval, LH = Limited hold, PRW = Premature response window.*

### Supplementary table 1

#### TABLE 1 | Stage parameters & requirements for the Visuospatial cue task

| Stage | Session (min) | ITI (s) | BSI1 (s) | CPT (s) | BSI2 (s) | PRW (s) | LH (s) | TO (s) | Stim | Criteria |
|---|---|---|---|---|---|---|---|---|---|---|
| H+C | 20 | - | - | - | - | - | - | - | - | 3 trials |
| 1 | 30 | 3 | - | - | - | - | - | - | - | Complete ≥80 trials |
| 2 | 30 | 3 | 0.5 | - | - | - | 30 | - | - | Complete ≥80 trials |
| 3 | 40 | 3 | 0.5 | 0.5 | 0.5 | 0.5 | 30 | 5 | - | ≥40 Correct trials |
| 4 | 40 | 3 | 0.5 | 0.5 | 0.5 | 0.5 | 30 | 5 | - | ≥40 Correct trials |
| 5.0 | 40 | 5 | 0.5 | 0.5 | 0.5 | 0.5 | 20 | 5 | - | ≥50 Correct trials |
| 5.1 | 40 | 5 | 1 | 0.5 | 1 | 0.5 | 10 | 5 | - | ≥50 Correct trials |
| 5.2 | 40 | 5 | 1.5 | 0.5 | 1 | 0.5 | 5 | 5 | - | ≥60% Accuracy<br>≤20% PR |
| 6 | 40 | 5 | 1.5 | 0.5 | 1 | 0.5 | 5 | 5 | Yes | ≥60% Accuracy<br>≥50 Response trials |
| 7 | 40 | 5 | 1.5 | 3.5/<br>4.5 | 1 | 3.5/<br>4.5 | 5 | 5 | Yes | ≥60% Accuracy<br>≥50 Response trials |
| 8 | 40 | 5 | 4 | 0.5 | 1 | 0.5 | 5 | 5 | Yes | ≥60% Accuracy<br>≥50 Response trials |
| 9 | 40 | 5 | 1.5 | 0.5 | 1 | 0.5 | 5 | 5 | Yes | ≥60% Accuracy<br>≥50 Response trials |

ITI = Inter trial interval, BSI = Black screen interval, CPT = Cue presentation time, PRW = premature response window, LH = Limited hold, TO = Timeout, Stim = Optogenetic Stimulation, H+C = Habituation + Conditioning, PR = Premature response, Om = Omissions. Criteria between H+C – Stage 5.2 are the criteria thresholds for moving to the next training stage, while the criteria given from Stage 6 – 9 are the session criteria required to be included in the data analysis.

### Supplementary videos

Ratarrest, VSCT_Stage 6, VSCT_Stage 7-9
